# Fine-tuning heat stress algorithms to optimise global predictions of mass coral bleaching

**DOI:** 10.1101/2021.04.14.439773

**Authors:** Liam Lachs, John C. Bythell, Holly K. East, Alastair J. Edwards, Peter J. Mumby, William J. Skirving, Blake L. Spady, James R. Guest

**Author notes:** Corresponding Author: Liam Lachs –.

## Abstract

Increasingly severe marine heatwaves under climate change threaten the persistence of many marine ecosystems. Mass coral bleaching events, caused by periods of anomalously warm sea surface temperatures (SST), have led to catastrophic levels of coral mortality globally. Remotely monitoring and forecasting such biotic responses to heat stress is key for effective marine ecosystem management. The Degree Heating Week (DHW) metric, designed to monitor coral bleaching risk, reflects the duration and intensity of heat stress events, and is computed by accumulating SST anomalies (HotSpot) relative to a stress threshold over a 12-week moving window. Despite significant improvements in the underlying SST datasets, corresponding revisions of the HotSpot threshold and accumulation window are still lacking. Here, we fine-tune the operational DHW algorithm to optimise coral bleaching predictions using the 5km satellite-based SSTs (CoralTemp v3.1) and a global coral bleaching dataset (37,871 observations, National Oceanic and Atmospheric Administration). After developing 234 test DHW algorithms with different combinations of HotSpot threshold and accumulation window, we compared their bleaching-prediction ability using spatiotemporal Bayesian hierarchical models and sensitivity-specificity analyses. Peak DHW performance was reached using HotSpot thresholds less than or equal to Maximum Monthly Mean SST and accumulation windows of 4 – 8 weeks. This new configuration correctly predicted up to an additional 310 bleaching observations compared to the operational DHW algorithm, an improved hit rate of 7.9 %. Given the detrimental impacts of marine heatwaves across ecosystems, heat stress algorithms could also be fine-tuned for other biological systems, improving scientific accuracy, and enabling ecosystem governance.

## Introduction

Anthropocene marine heatwaves are becoming increasingly intense, more frequent and longer lasting due to climate change (Oliver et al. 2018; Holbrook et al. 2019). These anomalous heat stress events can have severe implications for a range of marine biota, e.g., influencing shifts in zooplankton communities, declines in key groups such as krill (Jiménez-Quiroz et al. 2019; Evans et al. 2020; Işkın et al. 2020), die-offs and reproductive failures of sea-birds (Cavole et al. 2016; Jones et al. 2018; Piatt et al. 2020), marine mammal strandings (Cavole et al. 2016), and mass coral bleaching and mortality events (Hughes et al. 2018). While surveying in situ ecosystem responses to climate change disturbances are essential to assess impact, it is also very costly. Accurate monitoring of ecosystem stress remotely and at scale is therefore crucial for effectively managing marine ecosystems and accurately predicting the impacts of climate change on marine biota. While satellite-based remote monitoring and forecasting programmes have been implemented across various biological contexts, we focus this study specifically on remote monitoring and forecasting of coral bleaching. Coral reefs are highly productive ecosystems that provide habitat to over a million marine species and essential ecosystem services (e.g., coastal protection, food, fisheries and tourism livelihoods) to hundreds of millions of people, estimated to be worth over 350,000 USD/ha/yr globally (Costanza et al. 2014; Ferrario et al. 2014). These ecosystems are increasingly faced with mass coral bleaching and mortality events (Hughes et al. 2017). The process of coral bleaching involves a breakdown in the symbiosis between coral hosts and their endosymbiotic phototrophic algae, and can ultimately lead to full or partial colony mortality (Brown 1997) and sub-lethal effects such as reduced growth (Edmunds 2005). Coral bleaching is a stress response with a variety of triggers (e.g., 2003anomalous temperature, both high and low; anomalous increases in the level of light; anomalous levels of salinity, both high and low; reduction in water quality; and diseases; Skirving et al. 2018). Episodes of mass coral bleaching occur across large spatial scales, affect numerous coral taxa, and can destroy entire healthy reefs within months. Pantropical mass bleaching events are becoming recurrent and are caused by the widespread increasing incidence of marine heatwaves under climate change (Hughes et al. 2017; Donner et al. 2018; Hoegh-Guldberg et al. 2019).

Over the past two decades, the National Oceanic and Atmospheric Administration’s (NOAA) Coral Reef Watch (CRW) programme has developed a suite of tools for monitoring coral bleaching risk using satellite-based sea surface temperature (SST) products. Specifically, the Degree Heating Week (DHW) metric is used as an indicator of heat stress levels sufficient to induce coral bleaching. DHW is computed as the accumulation of positive temperature anomalies (HotSpot) above a hypothesised coral bleaching stress temperature (i.e., 1°C above the Maximum of Monthly Means SST climatology – MMM) over the previous 12 weeks (Liu et al. 2003; Skirving et al. 2020). The DHW algorithm was designed in the 1990s, and the HotSpot threshold of *1*°C *above MMM* and accumulation window of *12 weeks* were chosen based on field and experimental evidence from Panama and the Caribbean (Glynn and D’Croz 1990; Jokiel and Coles 1990). Reflecting the technological advancements in remote-sensing capabilities since then, the SST and DHW products have increased in spatial resolution (50 km to 5 km) and temporal resolution (twice weekly to daily) (Liu et al. 2014). Despite these improvements, there has not yet been a corresponding revision of the HotSpot threshold and accumulation window used in the operational DHW algorithm.

Alternate DHW algorithms have been applied to evaluate associations between heat stress and coral bleaching, mostly at local or regional scales (Weeks et al. 2008; Guest et al. 2012; Kim et al. 2019; McClanahan et al. 2020; Wyatt et al. 2020). Particularly for weak marine heatwaves associated with coral bleaching, computing DHWs with a lower HotSpot threshold has proven useful for monitoring bleaching impacts and severity (Guest et al. 2012; Kim et al. 2019; Wyatt et al. 2020). Evidence also suggests that using a shorter accumulation window in the DHW algorithm can improve coral bleaching predictions in some cases (DeCarlo 2020; McClanahan et al. 2020). An optimisation study in which numerous DHW algorithms are tested against a global coral bleaching dataset could provide the scientific basis necessary to revise the operational DHW metric. Recently, DeCarlo (2020) showed that altering the HotSpot threshold and accumulation window can improve global coral bleaching prediction skill, based on weather forecasting techniques that predict bleaching events (yes or no) depending on whether DHWs exceed a certain threshold or not. DeCarlo used DHWs computed from Optimum Interpolation SST (OI-SSTv2) and coral bleaching records from a summative dataset of 100 well-studied coral reefs (Hughes et al. 2018). However, there is a mismatch in spatial scale between these two datasets; the SST data was extracted from 0.25-degree grid cells, while the area extent of each reef in the bleaching dataset ranged from 2 km^2^ (Southwest Rocks, Australia, and St. Lucia, South Africa) to over 9000 km^2^ (Northern Great Barrier Reef, Australia). Accordingly, there are potential mismatches between DHW values and bleaching data in their study. As such, there is a pressing need to apply a more comprehensive DHW optimisation study to a global dataset of direct bleaching observations and DHWs derived from a higher resolution SST dataset.

To construct a global coral bleaching model based on environmental covariates, predictions should account for spatial and temporal dependencies. For example, corals in certain geographic regions are likely to respond to heat stress with higher levels of coral bleaching (e.g., areas influenced by the El Niño Southern Oscillation) (Howells et al. 2016; Romero-Torres et al. 2020) and are likely to change through time due to coral adaptation and assemblage turnover (Dziedzic et al. 2019; Gouezo et al. 2019). From a statistical standpoint, spatiotemporal uncertainties in the bleaching–environment relationship must be accounted for to ensure that bleaching predictions are not just artefacts of spatial or temporal patterns in unmeasured variables. A number of studies modelling coral bleaching globally as a function of environmental covariates have assumed that the uncertainty of this relationship is spatiotemporally constant (Safaie et al. 2018; DeCarlo 2020). This assumption is unlikely to be true for coral bleaching responses, given the potential for coral adaptation (Bay et al. 2017; Matz et al. 2018) and the extent to which post-disturbance turnover can alter the composition of the coral assemblage (Gouezo et al. 2019) and therefore its tendency to experience subsequent coral bleaching. To address the spatial issues (but not temporal), Sully et al. (2019) introduced a Bayesian mixed modelling approach that explicitly resolved spatial variability in the uncertainty of bleaching– environment relationships. This was achieved by treating ecoregion and site as hierarchical random effects, but this comes at the cost of slow run-time, an issue further compounded by implementing these models via Monte Carlo Markov Chains (MCMC) which run iteratively and slowly (Rue et al. 2009). Given these issues, such an approach would not be appropriate for a coral bleaching optimisation study that aims to test hundreds of DHW algorithms whilst also accounting for spatial and temporal dependencies, since such a study would require a prohibitively large amount of computing resources.

This study seeks to offer a potential revision to the operational NOAA DHW metric with a view to improving its ability to predict mass coral bleaching. This will require a suitable methodology that is robust to spatiotemporal correlated uncertainties and runs with reasonable computational speed. Here, we apply an alternative approach to modelling bleaching–environment relationships based on Integrated Nested Laplace Approximation (INLA), which explicitly solves spatial and temporal uncertainties with much greater computational speed than MCMC (Rue et al. 2009). We aim to optimise two DHW algorithm parameters, the HotSpot threshold (from MMM – 4 to + 4°C) and the accumulation window (from 2 to 52 weeks) to improve coral bleaching predictions globally whilst still addressing the common issue of spatial and temporal dependencies. We achieved this by combining recently developed Bayesian hierarchical modelling techniques using INLA with a streamlined parallel-computing workflow on a high-performance computing cluster called “The Rocket”. This allowed hundreds of spatiotemporal INLA models to be run in a short time frame (i.e., hours instead of weeks as would be the case using MCMC).

## Data & Methods

### Coral Bleaching Data

The optimisation study presented here was based on a global dataset of 37,871 bleaching survey records from published and unpublished scientific sources spanning from 1969 to 2017 (Spady et al. 2021). Bleaching estimates were quantified by a wide range of surveying methods, including aerial surveys, line-intercept transects, belt transects, quadrats, radius plots, rapid visual assessments (e.g., manta tows), ad hoc estimates, and interviews with stakeholders. Since data were collected by hundreds of observers globally over several decades, data collection protocols for these different general methods are not standardised.

The original dataset underwent four layers of filtering *a priori* to ensure its suitability this for analyses. 1) Data were first filtered for errors. This excluded observations that did not have a recorded month or year, as well as observations in which the coordinates provided did not correspond with a coral reef location (5,562 observations excluded). 2) Data were removed if the survey date fell outside the period of peak thermal exposure for that year. As, for the purpose of this study, we are only interested in coral bleaching that results from thermal stress (i.e., not bleaching due to cold-stress, nutrient enrichment etc.), instances of bleaching that cannot be linked to the period of peak thermal exposure may not accurately reflect the status of heat-induced bleaching for that year and location. We defined the period of peak thermal exposure as the month prior to the month of MMM up to three months after the month of MMM. For example, if the month of MMM was February for a certain location, only observations from January-May were included. Further, we ensured that the observation was not made before the date of maximum DHW in that year (19,292 observations excluded). 3) To account for different sampling protocols in records of percentage bleaching, we computed bleaching as a binary variable. Bleaching estimates were reported as means, ranges, or broad categories. First, we summarised these as representative minimum and maximum percentages. Then, the absence of ecologically significant bleaching was defined as having a maximum estimation of 10% bleaching or less, while the presence of ecologically significant bleaching was defined as having a minimum estimation of 20% or greater. Observations in which the maximum estimation exceeded 10% while the minimum estimation remained below 20% were filtered out to reduce the chance of misrepresenting bleaching and no-bleaching observations (Fig. S1) (1,452 observations excluded). 4) Lastly, to account for spatiotemporal patchiness *a priori*, we only retained years which had greater than 100 independent observations, had a qualitatively even global distribution, and were not temporally isolated (i.e., the proceeding years also needed to meet the previous two criteria). This resulted in removal of all data before 2003. Despite having 345 bleaching records in 2002, all data from this year were removed as over 80% of records were from the Great Barrier Reef region alone (1,185 observations excluded). The resulting dataset included 10,380 unique observations between 2003 and 2017, with >171 observations per year and sufficient spatial representation for each year.

Accumulated heat stress is considered to be the mechanism causing mass coral bleaching (Heron et al. 2016; Skirving et al. 2019), and marine heatwaves typically occur across hundreds to thousands of kilometres on spatial scales of weather-systems. The vast majority of bleaching observations in the dataset are associated with mass bleaching events, but despite our filtration process, some bleaching observations will inevitably result from small scale local heat stress and other non-heat related factors. Since the models presented in this study are based solely on large scale accumulated heat stress, the model predictions we present reflect the mechanism of mass coral bleaching which is referred to from here on.

### Temperature Data

Heat stress metrics were derived from a combination of CoralTemp v3.1 (Skirving et al. 2020), a gap-free global 5km daily SST dataset from 1985 until present, and corresponding 5km MMM climatology (Skirving et al. 2020). At each spatially referenced survey record, environmental data were extracted from the 5km grid cell encompassing that coordinate. These data consisted of a single MMM value and a time series of daily SST from the start of the pre-survey year until the end of the survey year.

For the operational DHW metric used by NOAA (DHW_op_), daily HotSpots were calculated as daily positive SST anomalies relative to MMM (1). Time series of daily DHW_op_ were then computed using the standard NOAA CRW method (2). HotSpots greater than 1°C were accumulated across a 12-week moving window (84 days inclusive), where *i* is the date and *n* is the earliest date of the accumulation window. Each daily HotSpot used in the summation is divided by seven a priori, such that

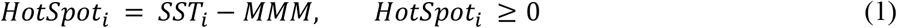

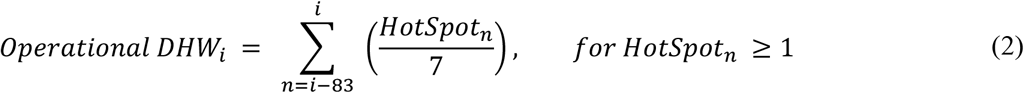

As an example, consider a 12-week window ending on April 1^st^ for a specific survey location. This window includes only three daily SSTs that exceed the MMM, equivalent to HotSpots of 0.5, 1.4, and 2.8°C. The DHW_op_ value for April 1st is the summation of 1.4 and 2.8°C each divided by seven, which is 0.6°C-weeks. The 0.5°C HotSpot value was not included in the summation as it was below 1°C (Skirving et al. 2020).

We computed a total of 234 test DHW metrics (DHW_test_), each with unique combinations of HotSpot thresholds (9 levels, from – 4 to + 4°C relative to MMM) and accumulation windows (26 levels, from 2 to 52 weeks). Unlike the operational metric, HotSpots for DHW_test_ were calculated relative to the MMM after an adjustment for the specific threshold in question (3). In the operational metric only HotSpots > 1°C are accumulated, however, in the test metrics all positive HotSpots are accumulated. Therefore, values of DHW_test_ are numerically different than DHW_op_ but are conceptually the same (see Figure 6). Time series of daily DHW_test_ were computed as the accumulation of HotSpots (4), where *i* is the date, *n* is the earliest date of the accumulation window, and *j* is the length of the accumulation window in days minus one, such that

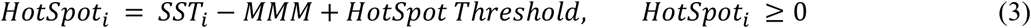

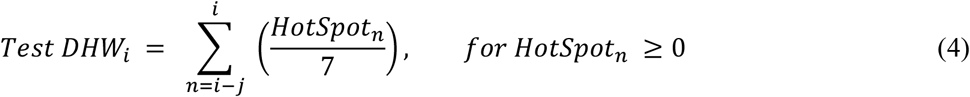

### Statistical Approach

The time unit used in the following models is the calendar year. As coral bleaching is more likely at higher levels of heat stress (Heron et al. 2016), the maximum of daily DHW values was computed from the year of each survey record. Thus, all further reference to DHW metrics relate to the annual maximum summary statistic. Given that the southern hemisphere summer starts before the end of the calendar year, there was a chance of misclassifying maximum DHW values. For instance, a maximum DHW on the first or last day of a calendar year will be part of the same heatwave event, however they will each be assigned to different calendar years. Previously, this has been addressed by adopting different calendars for each hemisphere (Skirving et al. 2019), however, this was not necessary in the current study since no such instances were present in the dataset. The relative performance of DHW metrics for predicting mass coral bleaching were assessed systematically using the following conceptual framework.

1. For each DHW metric, the association with coral bleaching was tested using a spatiotemporal Generalised Linear Model (GLM) with a Bernoulli error structure using INLA.
2. Sensitivity-specificity analysis was performed on this GLM to optimise predictions, tally model successes and failures, and provide metrics for model comparisons.
3. The first two steps were repeated for all DHW_test_ metrics and DHW_op_, resulting in 235 separate GLMs and sensitivity-specificity analyses, each run in parallel on separate Intel Xeon E5-2699 processors via the high-performance computing cluster “The Rocket”.
4. Model comparisons were used to determine the best-performing models and hence the optimal HotSpot threshold and accumulation window for predicting coral bleaching globally using DHWs.

### Model formulation

We have adopted a spatiotemporal Bayesian modelling approach to predict mass coral bleaching based on DHWs using the R-INLA package (http://www.r-inla.org) (Rue et al. 2009). Compared to more commonly used frequentist approaches, Bayesian inference allows uncertainty to be more easily interpreted. Moreover, using R-INLA over other Bayesian tools (e.g., Monte Carlo Markov Chains) provides the opportunity to resolve spatiotemporal correlation explicitly and more rapidly (Rue et al. 2009).

Observations of mass coral bleaching are often spatiotemporally correlated due to large-scale climatic drivers. While basic linear regressions applied to such data ignore these dependencies and lead to pseudoreplication (Hulbert 1984), R-INLA circumvents these issues. In each time point, spatial dependencies are dealt with by implementing the Matérn correlation across a Gaussian Markov random field (GMRF), essentially a map of spatially correlated uncertainty. This is achieved using stochastic partial differential equations (SPDE) solved on a Delaunay triangulation mesh of the study area. The parameters (*Ω*) that determine the Matérn correlation are the range (*r* – range at which spatial correlation diminishes) and error (*σ*). Weakly informative prior estimates of these parameters (*r*_*0*_ and *σ*_*0*_) are recommended when implementing the Matérn correlation (Fuglstad et al. 2019).

Temporal dependencies among these GMRFs are dealt with by imposing a first order autoregressive process (AR1), defined by the AR1 parameter (*ρ*) (9). This allows for correlation in model residuals through time avoiding pseudoreplication.

To test the effect of DHW metrics on coral bleaching, a triangular mesh (Fig. 2) was defined with a maximum triangle edge length of 600 km and a low-resolution convex hull (convex = -0.03) around the study sites to avoid boundary effects (1,790 nodes). This mesh was repeated for each year in the time series (26,400 nodes). The probability of coral bleaching for a given observation (*CB*_*t,i*_) in a given year (*t*) and location (*i*) was assumed to follow a Bernoulli distribution (π_t,i_) using the logit-link function for binary data. Bleaching was modelled as a function of the DHW metric in question (fixed effect: *DHW*_*t,i*_) whilst accounting for additional underlying spatiotemporal correlation among bleaching observations (random effect: *v*_*t,i*_),

**Figure 1.**
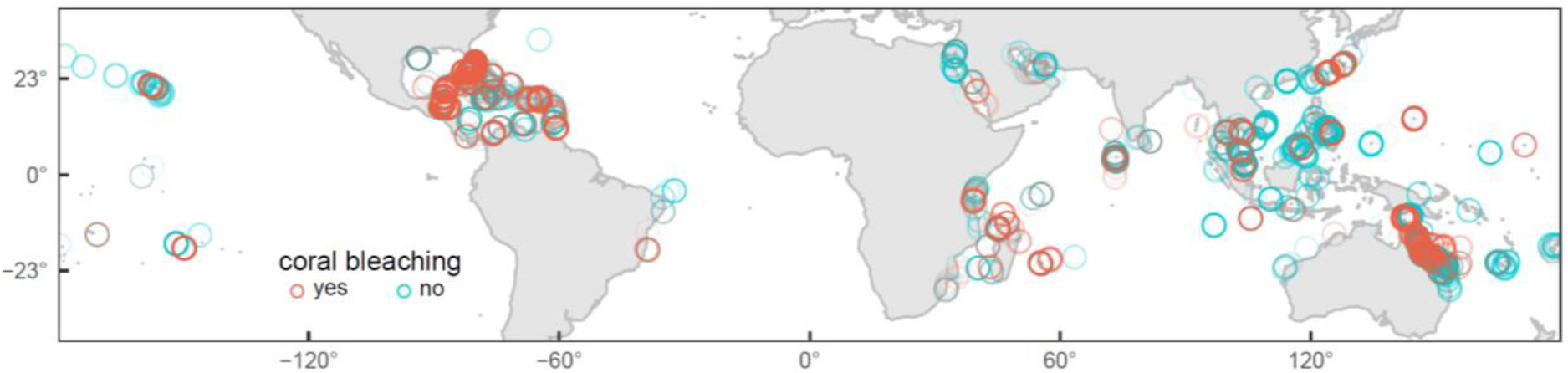
Distributions of coral bleaching survey records based on estimates of percentage coral bleaching (< 10% = no, > 20% = yes), measured at 5724 sites from 84 countries between 2003 and 2017 (N = 10,380) after four layers of *a priori* filtering (i.e., removal of errors, matching surveys with the period of peak thermal exposure in the year, accounting for inconsistent sampling protocols, and accounting for spatiotemporal patchiness).

**Figure 2.**
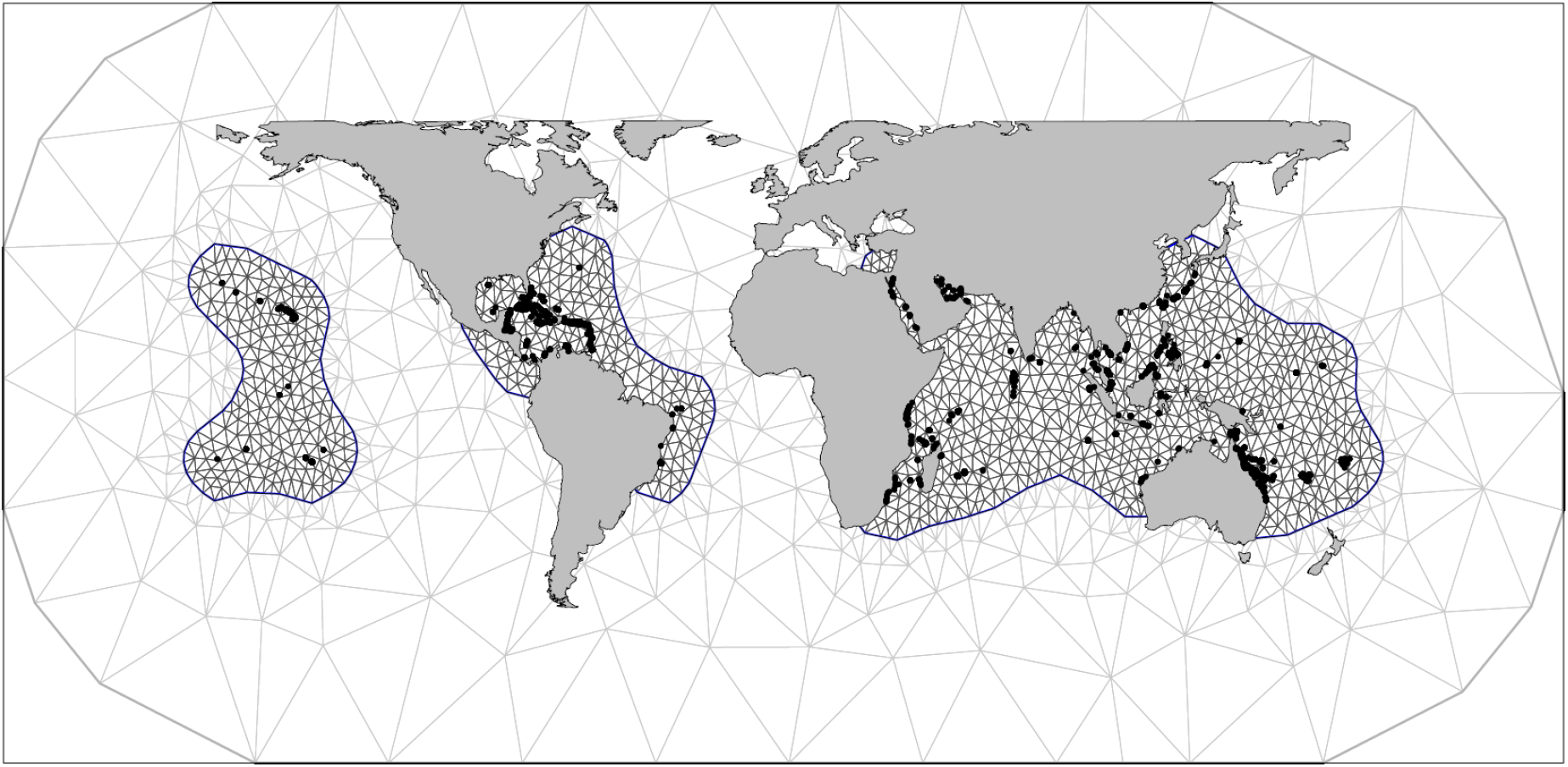
Constrained refined Delaunay triangulation mesh of 1790 nodes used for spatial correlation in one timestep. The spatiotemporal correlation over 15 years is computed over 15 such meshes totalling 26,400 nodes. Continents and bleaching survey coordinates (black points) overlay the higher resolution study area (black mesh) and lower resolution convex hull (grey mesh).

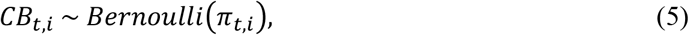

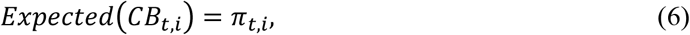

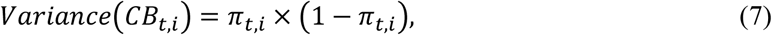

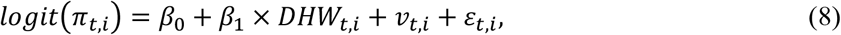

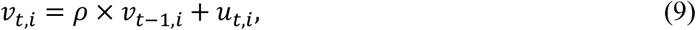

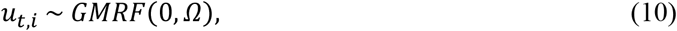

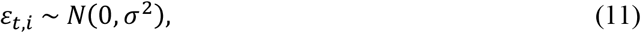

where *β*_*0*_ is the intercept, *β*_*1*_ is the DHW parameter estimate, *ρ* is the AR1 parameter, *u*_*t,i*_ represents the smoothed spatial effect from the GMRF mesh, elements of *Ω* (*r* and *σ*) are estimated from the Matérn correlation, and *ε*_*t,i*_ contains the independently distributed residuals. Following the recommendations from Fuglstad et al. (2019), we specified weakly informative priors for *r*_*0*_ (2000 km) and *σ*_*0*_ (1.15) based on the residual variogram and error from an intercept-only null Bernoulli GLM (Fig. S2). We also tested different priors; however, they had a negligible effect on the estimates of any model parameters. To avoid imposing artificial temporal dependencies, we used a non-informative default prior for *ρ*.

### Model Validation

Standard model validation steps were conducted for the best performing GLM and included plotting bleaching observations against fitted values, assessing model residuals for spatiotemporal correlation using maps and variograms, and producing a time series of maps showing spatiotemporally correlated uncertainty (Zuur and Ieno 2017). The dataset presented here was considerably patchy in both space and time despite prior filtering (e.g., no South Pacific observations in 2003, 2012, or 2013). Patchy data is a pertinent issue in statistics (Little and Rubin 2002) and can have a considerable effect on estimated model parameters (Bihrmann and Ersbøll 2015), and model selection criteria (e.g., Deviance Information Criterion – DIC) (Nakagawa and Freckleton 2008). Thus, to address patchiness beyond basic filtering, we performed a simulation test (Fig. S3 & Fig. S4). In summary, patchiness did not have an important effect on estimated model parameters (Fig. S5), validating the broader model comparison methods and results of the main study. Full details are described in the Supplementary Materials.

### Sensitivity-Specificity Analysis

To optimise binary predictions from each Bernoulli GLM, sensitivity-specificity analyses were performed using receiver operating characteristic (ROC) curves in R (Robin et al. 2011) without considering spatiotemporal dependencies. This method is commonly applied in bioinformatics and medical decision making to determine the performance of binary classifications. Here, sensitivity is defined as the proportion of correctly classified bleaching observations (true positives), and specificity as the proportion of correctly classified no-bleaching observations (true negatives). As a probability cut-off is moved over all possible values, the ROC plot shows the corresponding sensitivity and specificity at each level. The Area Under the Curve (AUC) from each ROC plot reflects the performance of that GLM relative to the perfect predictor (AUC = 1) and can be used for multi-model comparisons based on 95% confidence intervals computed using stratified bootstrap resampling (Robin et al. 2011). The hit rate, defined as the proportion of observed bleaching events that were correctly predicted, was also computed at the optimal cut-off level for each model.

### Model Comparisons

Model comparisons were based on the Bayesian DIC and two key metrics from the sensitivity-specificity analysis: AUC and hit rate. DIC is a measure of overall model parsimony (Zuur and Ieno 2017), but is based on both the DHW fixed effect and the spatiotemporal random effect. Therefore, AUC and bootstrapped confidence intervals were used as the preferred model comparison metric, as this evaluates the overall performance of a binary classifier relative to a perfect predicting model (Robin et al. 2011), based on the fixed effect only. Hit rate is an additional metric that allows easy interpretation of model success.

## Results

### Model Comparisons

For predicting coral bleaching based on DHW_test_, we identify (1) a group of worst performing models, (2) a group of better performing models, and (3) a suite of best performing models. (1) Poor GLM performance was associated with DHW_test_ metrics computed on HotSpot thresholds ≥ MMM + 2°C or accumulation windows ≥ 22 weeks. This was evident by low AUC values < 0.7 and high DIC values > 7000 (Fig. 3, right and upper regions). (2) The remaining GLMs (HotSpot threshold ≤ MMM + 1°C, accumulation window ≤ 20 weeks) were associated with better coral bleaching predictions (AUC) and model parsimony (DIC) (Fig. 3, lower and lower left regions). (3) Finer determination of the best models of this subset was made possible by incorporating sensitivity-specificity uncertainty into model comparisons (Fig. 4, 95% bootstrapped confidence intervals). A performance-optima relationship was apparent between AUC and the HotSpot threshold and accumulation window, whereby peak GLM performance was reached when DHW accumulation windows were 4 – 8 weeks (Fig. 4). When DHW accumulation windows were outside this range (2 weeks or ≥ 10 weeks), corresponding AUC was significantly lower than the AUC of the best performing GLMs (Fig. 4, blue shaded region). Notably, of all the GLMs that used the same accumulation window (grey and white band groupings, Fig. 4), those models applying lower HotSpot thresholds performed better in terms of AUC and DIC. The 8-week accumulation window resulted in the best overall fit of AUC and DIC combined (max DIC = 6812). In summary, the suite of best-performing models (in terms of bleaching prediction) applied DHW_test_ metrics based on HotSpot thresholds ≤ MMM and accumulation windows of 4 – 8 weeks.

**Figure 3.**
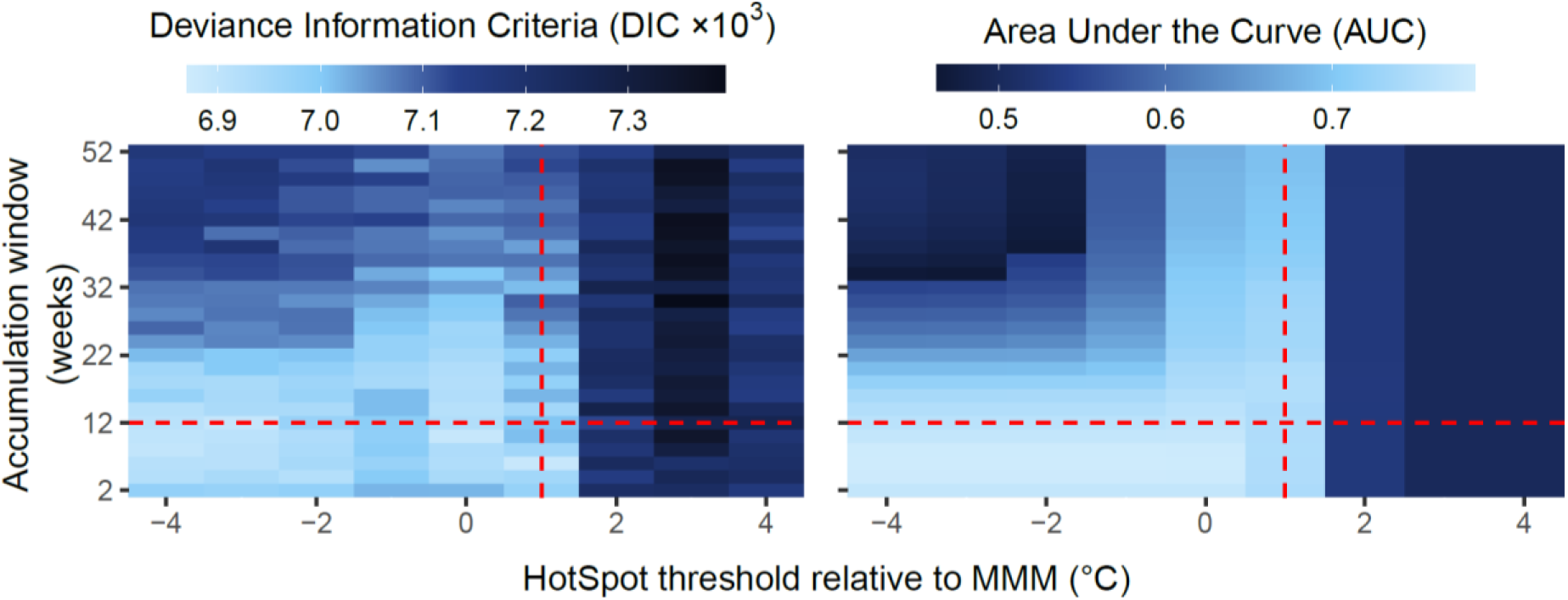
Model comparison heatmaps showing the Deviance Information Criterion (DIC) and Area Under the Curve (AUC) for 234 Generalised Linear Models (GLMs) that each predict coral bleaching based on a different DHW_test_ metric. Raster cells represent individual GLMs plotted by HotSpot threshold and accumulation window. The threshold and window used for DHW_op_ are shown by red dashed lines (MMM + 1°C, and 12-weeks). Results for the DHW_op_ GLM are not shown on the heat maps (DIC = 6967, AUC = 0.758).

**Figure 4.**
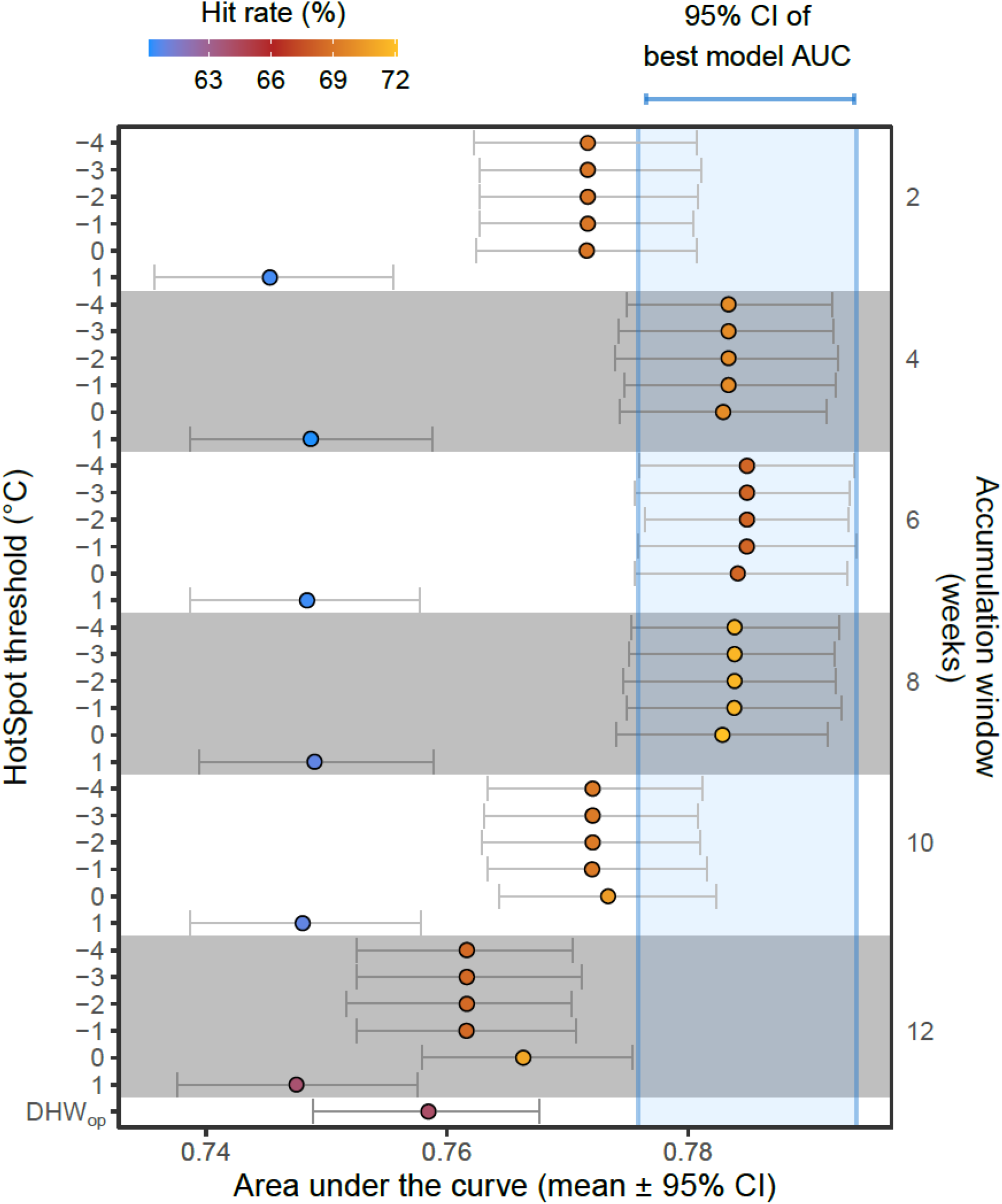
Model comparisons accounting for uncertainty in Area Under the Curve (AUC) showing the mean and 95% bootstrapped confidence intervals (CI). Each point represents a Generalised Linear Model GLM that predicts coral bleaching based on a different DHW_test_ metric, ordered by HotSpot threshold and accumulation window (both increasing downwards). The hit rate (proportion of observed bleaching events correctly predicted) is shown for each GLM (point colour) and the AUC of the best GLM is shown as a blue shaded region. Note the DHW_op_ algorithm is slightly different than the DHW_test_ algorithm (Equation 1-4).

### Best Model – Validation

The GLM based on the DHW_test_ metric with HotSpot threshold of MMM + 0°C and accumulation window of 8 weeks (DHW_test-0C-8wk_), was a representative of the suite of best-performing models. The probability of bleaching output from this model (based on DHW_test-0C-8wk_ and unmeasured spatiotemporally correlated factors) closely matched the observational bleaching record (Fig. 5A). Both the fixed effect (DHW_test-0C-8wk_) and the random effect (spatiotemporal uncertainty) provided important contributions to predictions of coral bleaching (Fig. S7). The sensitivity-specificity analysis reflected the high performance for this model, with an AUC value of 0.783 (Fig. 5B). The range parameter (*r*) of GMRFs showed that drivers of bleaching other than DHW_test-0C-8wk_ were spatially correlated up to 697 km (Fig. S6), consistent with the spatial scale of climatic and weather systems. The AR1 parameter (*ρ*) of 0.62 indicated moderate temporal correlation of uncertainty in predicted coral bleaching (i.e., drivers other than DHW_test-0C-8wk_), meaning that the uncertainty in bleaching predictions in one year is affected by that of the previous year by a factor of 0.62 (Fig. S6). This can be seen visually on maps of temporally correlated GMRFs (Fig. S7).

**Figure 5.**
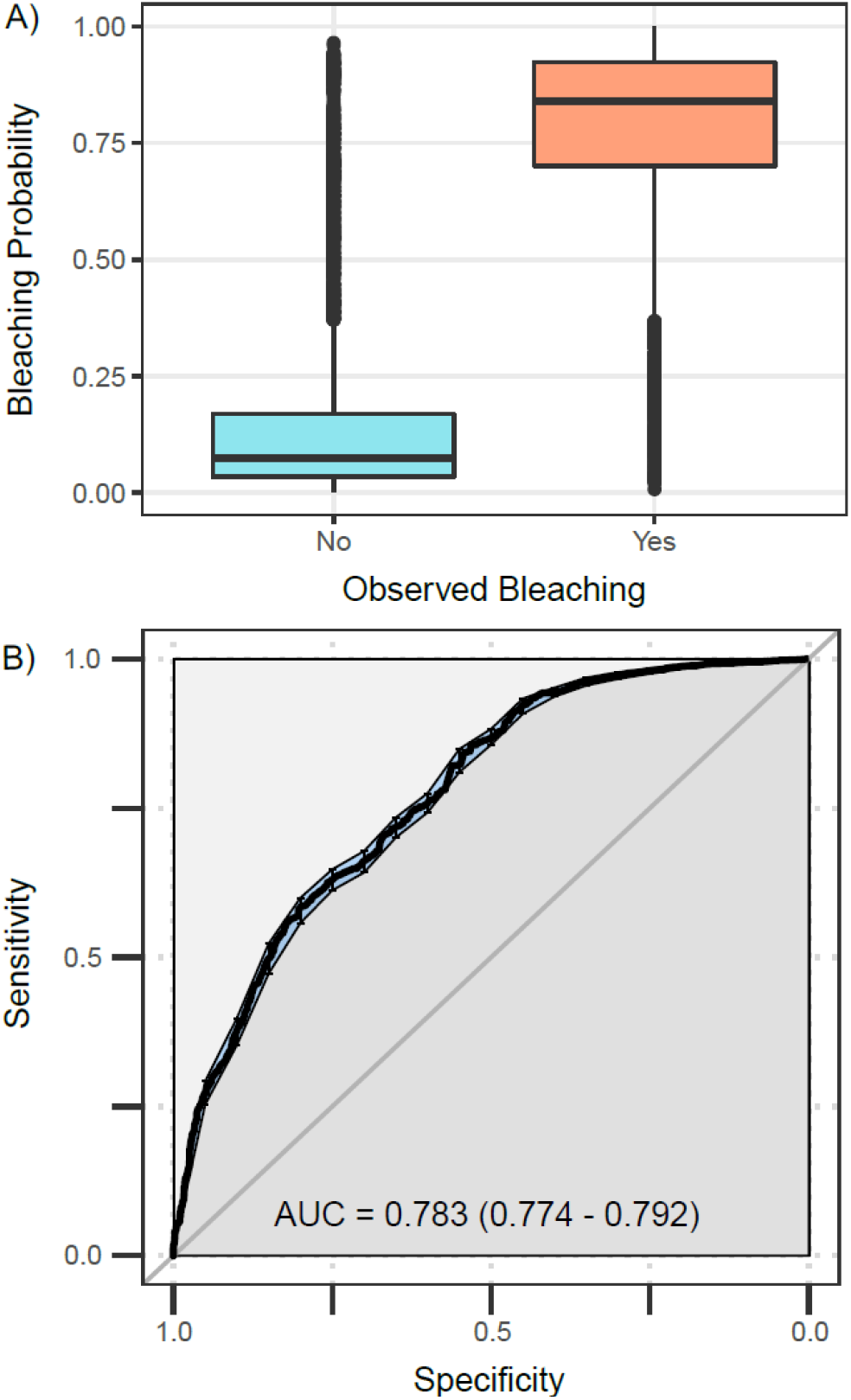
Exploration of best-performing GLM which predicts coral bleaching based on DHW_test-0C-8wk_ (HotSpot threshold = MMM, accumulation window = 8 weeks) and spatiotemporal uncertainty. (A) Fitted values or bleaching probabilities are shown relative to bleaching observations from the global dataset, showing a clear separation between bleaching and non-bleaching categories. (B) Sensitivity-specificity analysis is shown for the same GLM without spatiotemporal uncertainty. Sensitivity is defined as the proportion of correctly classified bleaching observations (true positives), and specificity as the proportion of correctly classified no-bleaching observations (true negatives).

Area Under the Curve (AUC) and bootstrapped 95% confidence intervals (shown in brackets) reflect the distance to a perfect predicting model (AUC = 1).

### Best Model – Understanding Heat Stress

Even though lowering the HotSpot threshold and reducing the accumulation window improved predictions of mass coral bleaching (Fig. 3, Fig. 4), the DHW_op_ metric still categorised bleaching observations well. DHW_op_ values were greater for bleaching records than for non-bleaching records (Fig. 6). Of the 517 highest heat stress records (> 95^th^ percentile: > 9.0°C-weeks), 78% were associated with coral bleaching observations, highlighting the importance of heat stress as a proximate cause of coral bleaching. Such levels of heat stress relate to NOAA CRW Bleaching Alert Level 2. However, in comparison to DHW_op_, the test metric DHW_test-0C-8wk_ showed a higher distribution of heat stress values overall, but lower extremes values (Fig. 6). This is due to a lower HotSpot threshold and shorter accumulation window, respectively. This was characterised by fewer DHW values of zero (1 vs. 27%), a higher mean (5.2 vs. 2.5°C-weeks), a higher 95^th^ percentile (9.9 vs. 9.0°C-weeks), but a lower 99^th^ percentile (11.3 vs. 12.5°C-weeks). The number of bleaching observations associated with a heat stress of zero was 6 for DHW_test-0C-8wk_ and 122 for DHW_op_. Given that DHW_test-0C-8wk_ had a lower HotSpot threshold, fewer bleaching observations are associated with heat stress values of zero. In other words, reducing the HotSpot threshold increased our ability to predict coral bleaching associated with weak marine heatwaves.

**Figure 6.**
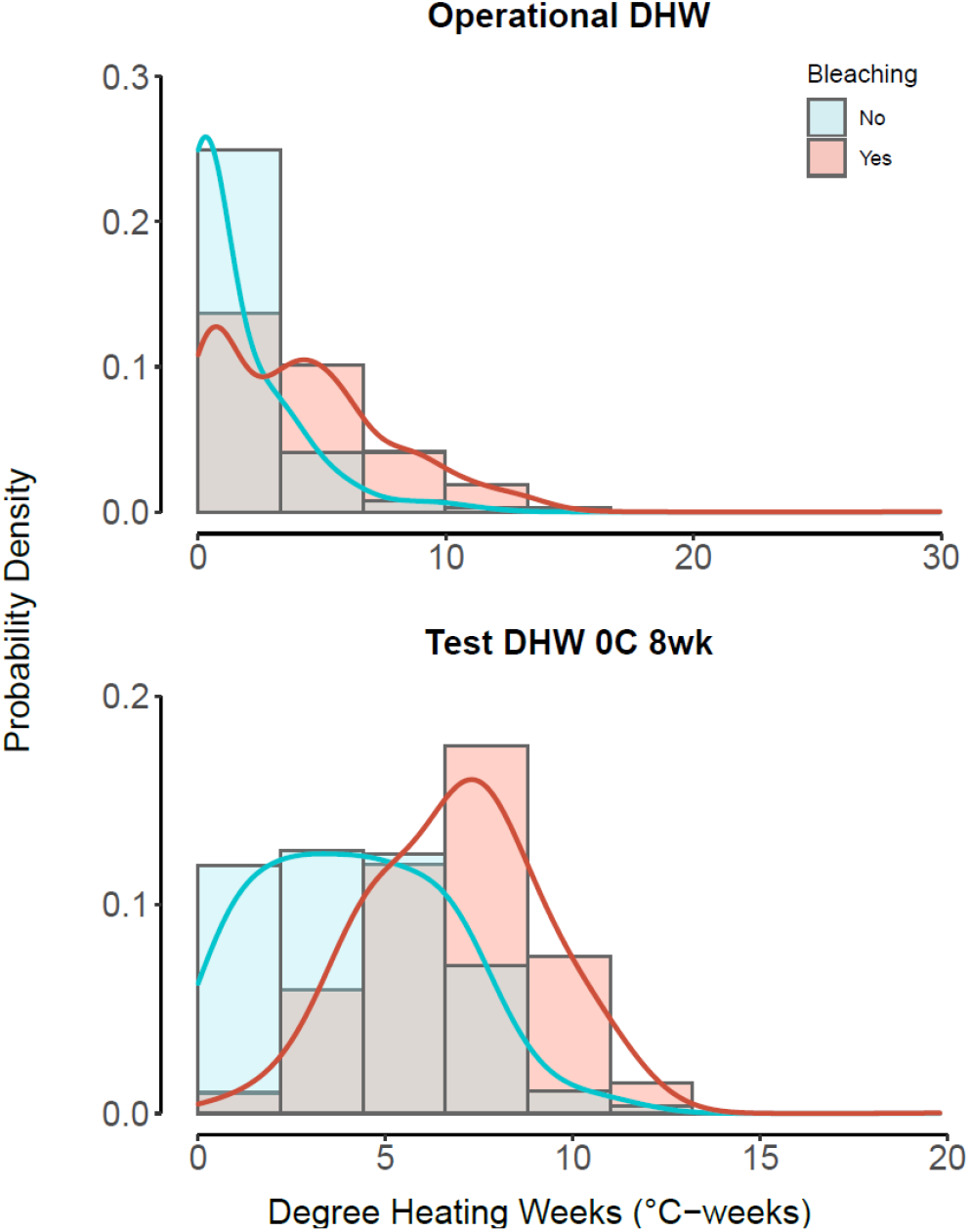
DHW distributions for bleaching records (red) and non-bleaching records (blue), shown as histograms and probability density curves. For comparison of different DHW metrics, the operational metric used by NOAA (DHW_op_) is shown alongside one of the best-performing metrics (DHW_test-0C-8wk_), calculated using a lower HotSpot threshold (MMM + 0°C) and a smaller accumulation window (8 weeks).

## Discussion

Heat stress can have considerable impacts on marine organisms and entire marine ecosystems (Eakin et al. 2019; Smale et al. 2019). The DHW metric is a measure of accumulated heat stress widely used to predict mass coral bleaching caused by anomalous temperatures above typical summertime conditions (Heron et al. 2016; Safaie et al. 2018; Skirving et al. 2019; Sully et al. 2019). The remote-sensed SST products underpinning the operational NOAA DHW metric have improved stepwise over the last two decades (Wellington et al. 2001; Liu et al. 2003; Liu et al. 2013; Skirving et al. 2020), however, there has not yet been a corresponding revision of the HotSpot threshold and accumulation window used in this algorithm. Here, we developed 234 different DHW algorithm variants each with a different HotSpot threshold and accumulation window. We assess the performance of these DHW_test_ metrics for predicting mass coral bleaching globally. Compared to DHW_op_, it was possible to improve the coral bleaching hit-rate by up to 7.9% by using different HotSpot thresholds and accumulation windows, equating to an additional 310 correctly predicted bleached reefs out of a total of 3895 (also linked to an increased false negative rate of 3%). Simply reducing the HotSpot threshold to MMM (or < MMM) rather than MMM + 1°C, resulted in up to 6.8% increases in hit rate, whilst using an accumulation window of 8 weeks instead of 12 weeks maximised this hit rate. Such improvements were also reflected in model comparison metrics from sensitivity-specificity analyses (increased AUC of 0.02) and Bayesian inference (decreased DIC of 36). Models using the 4 – 8 week accumulation window generally performed best, reflecting the typical duration of the vast majority of coral bleaching heat stress events to date (Oliver et al. 2018). Under climate change, however, average sea temperatures and the duration of marine heatwaves are predicted to continue increasing (Hoegh-Guldberg et al. 2018; Oliver et al. 2018), meaning in the future, longer DHW accumulation windows may better capture the levels of heat stress relevant to coral bleaching. Given that baselines are shifting throughout biotic and abiotic marine systems and that rates of adaptation to future environmental conditions are yet unknown, the concepts addressed in this study likely need to be revisited in the future at semi-regular intervals to ensure that the DHW product remains as accurate as possible.

### Complexities of coral bleaching

Coral bleaching is a stress response whereby photosynthetic algal symbionts are lost from the coral host tissues, resulting in the white coral skeleton becoming progressively more visible (Brown 1997; Douglas 2003). Given the complexity of this host-symbiont relationship, survey metrics such as ‘coral bleaching extent’ provide limited information from which to infer biological causes. Coral bleaching is affected by numerous biological factors including symbiont community composition and their environmental responses (e.g., more or less heat-tolerant algal taxa) (LaJeunesse et al. 2018), host heterotrophy (e.g., reliance on the symbiont) (Conti-Jerpe et al. 2020), the capacity for acclimation and adaptation both genetic and epigenetic (intra- and inter-generational) (Kirk et al. 2018; Liew et al. 2020), and coral taxonomy (e.g., different life history strategies) (Marshall and Baird 2000; Guest et al. 2012). In addition, other environmental factors can influence bleaching responses in corals, such as high solar insolation, cloudiness, winds, tidal extremes, thermal variability, cold-water stress and nutrient enrichment (Mumby et al. 2001; Hoegh-Guldberg et al. 2005; Anthony et al. 2007; Anthony and Kerswell 2007; Wiedenmann et al. 2013; González-Espinosa and Donner 2020). Given this suite of biotic and abiotic factors, a perfect-predicting coral bleaching algorithm would need to combine heat-stress metrics with other environmental and biological parameters that in many cases are often not available. NOAA CRW are investigating the potential improvements to DHW via the inclusion of solar insolation with the development of their Light Stress Damage (LSD) satellite-based product (Skirving et al. 2018).

Here we have refined the ability of a common heat stress metric to predict mass coral bleaching. Ideally, such an optimisation study would be based on coral bleaching data that relate to only heat stress related mechanisms. By filtering the dataset as described, we did our best to achieve this, however, bleaching observations in the dataset may inevitably have been caused by other biotic or abiotic factors, contributing to the noise in our results. Bleaching observations from surveys may also be subject to other inaccuracies such as the assumption that sampling only part of a reef is representative of the entire reef. Despite these points, the model comparisons performed in this study remain valid as model biases were applied to all models equally. Given these facts, the AUC and hit rate from sensitivity-specificity analyses are unlikely to reflect the absolute accuracy of DHW metrics, but rather allow comparisons of relative accuracy to determine optimal HotSpot thresholds and accumulation windows. The optimisation study presented here was performed on a global coral bleaching dataset. For scientists and practitioners aiming to assess global patterns in coral bleaching, we have shown that bleaching predictions can be improved by computing DHW metrics using an optimal HotSpot threshold of the MMM + 0°C and accumulation window of 8 weeks. These recommended DHW algorithm refinements are only applicable to global analyses and predictions of mass coral bleaching caused by heat stress. Moreover, it is important to note that the quasi-opportunistic nature of coral bleaching surveys (i.e., monitoring coral bleaching when DHW values are high indicating high bleaching risk) can lead to a confirmation bias in studies of coral bleaching and heat stress. Monitoring programmes should address this limitation, by aiming to survey bleaching more regularly, even when there is no accumulated temperature stress (i.e., DHW = 0).

### Global and regional scales

A regionally sensitive DHW algorithm would likely improve predictions of mass coral bleaching. For instance, many scientific studies have used variants of the DHW algorithm to better predict coral bleaching in their study site (Guest et al. 2012; Kim et al. 2019; Wyatt et al. 2019). This will likely continue, since oceanographic and climatic systems, coral assemblages, and the distribution of algal symbiont taxa vary geographically and at regional scales (Veron 1995; Clarke 2014; LaJeunesse et al. 2018). For instance, the thermal regime of the tropical Eastern Pacific is distinct from many other tropical regions, characterised by high variability due to the El Niño Southern Oscillation, with more intense warm water conditions typical of La Niña years compared to El Niño years (Clarke 2014). Long-term trends in coral coverage from this region, which have remained very stable over the past 3 decades, are atypical compared to most tropical reefs which have suffered persistent declines (Hughes et al. 2017; Romero-Torres et al. 2020). Such distinct trends in the tropical Eastern Pacific could be caused by adaption of corals there to highly variable thermal regimes (Romero-Torres et al. 2020). This is just one example of a region that could benefit from a specific regional DHW optimisation. Notably, the methods applied in this study would be easily adapted to develop such regional DHW products.

### Future outlook

Optimising heat stress metrics for specific purposes could also be useful for other marine systems. Marine heatwaves have contributed to marked ecological disturbances beyond mass coral bleaching and mortality events (Ummenhofer and Meehl 2017; Frölicher and Laufkötter 2018; Smale et al. 2019), yet specific metrics to predict these other disturbances are not often implemented. The northeast Pacific warming event of 2013 – 2015, termed “the blob”, was the subject of unusually high SST anomalies and repeated marine heatwaves (Di Lorenzo and Mantua 2016). The blob was associated with considerable ecological impacts, including the mass stranding of marine mammals such as sea lion and whales (Cavole et al. 2016), die-offs and reproductive failure of seabird populations (Cavole et al. 2016; Jones et al. 2018; Piatt et al. 2020), and reduced survival and growth of foraging fish (von Biela et al. 2019). In all these cases, evidence suggested that declines in higher trophic levels were associated not to direct effects of heat stress, but to the cascading effects of heat-mediated declines at lower trophic levels. Reduced abundance and altered composition of zooplankton communities including krill are highly susceptible to heat stress (Jiménez-Quiroz et al. 2019; Evans et al. 2020; Işkın et al. 2020), which can result in reduced food availability for higher trophic level animals (e.g., Cassin’s auklet and Californian sea lion), their emaciation and mortality (Cavole et al. 2016). The urgency to understand the full extent of ecological impacts associated with marine heatwaves could in part be addressed by creating new heat stress indicators that are optimised for specific disturbances using similar methods to those applied here. While this would not allow for rapid response actions to such events, it would guide marine protected area design (i.e., focus on conserving thermal refugia) and inform future projections of marine systems and related policy recommendations.

### Conclusion

The Anthropocene is characterised by shifting baselines of biological communities, loss of biodiversity, and increasingly severe and frequent climatic disturbances. Thus, there is growing need to understand and be able to predict climatic and anthropogenic disturbances on habitats, particularly those that provide key ecosystem services to socioecological systems. Here, we have fine-tuned a commonly used heat stress algorithm to a specific purpose (i.e., predicting mass coral bleaching), and have shown that simple changes (compared to the operational algorithm) can result in a considerable improvement in prediction success. The philosophy behind this optimisation study was to remove prior expectations, run the models, and allow the data to reveal the optimal HotSpot threshold and accumulation window for predicting mass coral bleaching globally. In this case, coral bleaching observations were correctly predicted up to 7.9% more often just by reducing the HotSpot threshold and accumulation window of the DHW_test_ metric. Broadly, improving bleaching prediction success of the operational DHW metric can support stakeholders and end-users such as coral reef managers, inform the design of MPA networks (e.g., including thermal refugia), and provide more accurate information which can lead to better conservation and restoration decision-making (e.g., shifting valuable coral nurseries during heatwaves, assisting with decisions on when to relocate acquarium-grown corals to the reef, etc.). Fine-tuning DHWs also has potential for other specific systems, such as predicting planktonic shifts and associated impacts to higher trophic levels. Increasingly under climate change, marine heatwaves are shaping species populations, biological food webs and even ecosystem structure and function (Hughes et al. 2017; Eakin et al. 2019; Smale et al. 2019). Thus, optimising our predictions of heat stress and the associated ecological impacts will be key to understanding the future of marine ecosystems.

## Acknowledgements

This research was funded by the Natural Environment Research Council’s ONE Planet Doctoral Training Partnership (NE/S007512/1) to L.L., the European Research Council Horizon 2020 project CORALASSIST (Project number 725848) to J.R.G. and A.J.E. Coral Reef Watch and ReefSense staff (B.L.S. and W.J.S.) were supported by NOAA grant NA19NES4320002 (Cooperative Institute for Satellite Earth System Studies) at the University of Maryland/ESSIC, and the U.S. Department of Defense’s Strategic Environmental Research and Development Program. The authors would also like to thank Dr. Adriana Humanes and Prof. Stephen Rushton for their intellectual contributions to the study and methodology, and the innumerable divers, volunteers and citizen scientists who helped with coral bleaching data collection. The scientific results and conclusions, as well as any views or opinions expressed herein, are those of the author(s) and do not necessarily reflect the views of NOAA or the Department of Commerce.

## Author Contributions

L.L., J.C.B., A.J.E., J.R.G., and W.J.S. conceived and designed the study; the National Oceanic and Atmospheric Administration Coral Reef Watch program provided the coral bleaching data and sea surface temperature data; B.L.S., L.L., and W.J.S. accessed and filtered the datasets; L.L. developed the models, prepared the figures, and wrote the code; L.L., and B.L.S. wrote the first draft of the paper; and J.C.B., H.K.E., A.J.E., J.R.G., P.J.M., and W.J.S. contributed significantly to the interpretation and editing of the manuscript.

## References

Anthony KRN, Connolly SR, Hoegh-Guldberg O (2007) Bleaching, energetics, and coral mortality risk: Effects of temperature, light, and sediment regime. Limnol Oceanogr 52:716–726. https://doi.org/10.4319/lo.2007.52.2.0716

Anthony KRN, Kerswell AP (2007) Coral mortality following extreme low tides and high solar radiation. Mar Biol 151:1623–1631. https://doi.org/10.1007/s00227-006-0573-0

Bay RA, Rose NH, Logan CA, Palumbi SR (2017) Genomic models predict successful coral adaptation if future ocean warming rates are reduced. Sci Adv 3:1–10. https://doi.org/10.1126/sciadv.1701413

Bihrmann K, Ersbøll AK (2015) Estimating range of influence in case of missing spatial data: A simulation study on binary data. Int J Health Geogr 14:p. https://doi.org/10.1186/1476-072X-14-1

Brown BE (1997) Coral bleaching: Causes and consequences. Coral Reefs 16:. https://doi.org/10.1007/s003380050249

Cavole LM, Demko AM, Diner RE, Giddings A, Koester I, Pagniello CMLS, Paulsen ML, Ramirez-Valdez A, Schwenck SM, Yen NK, Zill ME, Franks PJS (2016) Biological impacts of the 2013– 2015 warm-water anomaly in the northeast Pacific: Winners, Losers, and the Future. Oceanography 29:273–285. https://doi.org/10.5670/oceanog.2016.32

Clarke AJ (2014) El Niño Physics and El Niño Predictability. Ann Rev Mar Sci 6:79–99. https://doi.org/10.1146/annurev-marine-010213-135026

Conti-Jerpe IE, Thompson PD, Wong CWM, Oliveira NL, Duprey NN, Moynihan MA, Baker DM (2020) Trophic strategy and bleaching resistance in reef-building corals. Sci Adv 6:p. https://doi.org/10.1126/sciadv.aaz5443

Costanza R, de Groot R, Sutton P, van der Ploeg S, Anderson SJ, Kubiszewski I, Farber S, Turner RK (2014) Changes in the global value of ecosystem services. Glob Environ Chang 26:152–158. https://doi.org/10.1016/j.gloenvcha.2014.04.002

DeCarlo TM (2020) Treating coral bleaching as weather: a framework to validate and optimize prediction skill. PeerJ 8:e9449. https://doi.org/10.7717/peerj.9449

Di Lorenzo E, Mantua N (2016) Multi-year persistence of the 2014/15 North Pacific marine heatwave. Nat Clim Chang 6:1042–1047. https://doi.org/10.1038/nclimate3082

Donner SD, Heron SF, Skirving WJ (2018) Future Scenarios: A Review of Modelling Efforts to Predict the Future of Coral Reefs in an Era of Climate Change. In: Oppen MJH van, Lough JM (eds) Coral Bleaching, Ecological Studies 233, Second Edi. Springer International Publishing AG, part of Springer Nature 2018, pp 325–341

Douglas AE (2003) Coral bleaching - How and why? Mar Pollut Bull 46:385–392. https://doi.org/10.1016/S0025-326X(03)00037-7

Dziedzic KE, Elder H, Tavalire H, Meyer E (2019) Heritable variation in bleaching responses and its functional genomic basis in reef-building corals (Orbicella faveolata). Mol Ecol 28:2238–2253. https://doi.org/10.1111/mec.15081

Eakin CM, Sweatman HPA, Brainard RE (2019) The 2014–2017 global-scale coral bleaching event: insights and impacts. Coral Reefs 38:539–545. https://doi.org/10.1007/s00338-019-01844-2

Edmunds PJ (2005) The effect of sub-lethal increases in temperature on the growth and population trajectories of three scleractinian corals on the southern Great Barrier Reef. Oecologia 146:350– 364. https://doi.org/10.1007/s00442-005-0210-5

Evans R, Lea MA, Hindell MA, Swadling KM (2020) Significant shifts in coastal zooplankton populations through the 2015/16 Tasman Sea marine heatwave. Estuar Coast Shelf Sci 235:106538. https://doi.org/10.1016/j.ecss.2019.106538

Ferrario F, Beck MW, Storlazzi CD, Micheli F, Shepard CC, Airoldi L (2014) The effectiveness of coral reefs for coastal hazard risk reduction and adaptation. Nat Commun 5:1–9. https://doi.org/10.1038/ncomms4794

Frölicher TL, Laufkötter C (2018) Emerging risks from marine heat waves. Nat Commun 9:2015– 2018. https://doi.org/10.1038/s41467-018-03163-6

Fuglstad GA, Simpson D, Lindgren F, Rue H (2019) Constructing Priors that Penalize the Complexity of Gaussian Random Fields. J Am Stat Assoc 114:445–452. https://doi.org/10.1080/01621459.2017.1415907

Glynn PW, D’Croz L (1990) Experimental evidence for high temperature stress as the cause of El Niño-coincident coral mortality. Coral Reefs 8:181–191. https://doi.org/10.1007/BF00265009

González-Espinosa P, Donner S (2020) Cold-water coral bleaching prediction: role of temperature, and potential integration of light exposure. Mar Ecol Prog Ser 642:133–146. https://doi.org/10.3354/meps13336

Gouezo M, Golbuu Y, Fabricius K, Olsudong D, Mereb G, Nestor V, Wolanski E, Harrison P, Doropoulos C (2019) Drivers of recovery and reassembly of coral reef communities. Proc R Soc B Biol Sci 286:. https://doi.org/10.1098/rspb.2018.2908

Guest JR, Baird AH, Maynard JA, Muttaqin E, Edwards AJ, Campbell SJ, Yewdall K, Affendi YA, Chou LM (2012) Contrasting patterns of coral bleaching susceptibility in 2010 suggest an adaptive response to thermal stress. PLoS One 7:1–8. https://doi.org/10.1371/journal.pone.0033353

Heron SF, Johnston L, Liu G, Geiger EF, Maynard JA, De La Cour JL, Johnson S, Okano R, Benavente D, Burgess TFR, Iguel J, Perez DI, Skirving WJ, Strong AE, Tirak K, Eakin CM (2016) Validation of Reef-Scale Thermal Stress Satellite Products for Coral Bleaching Monitoring. Remote Sens 8:

Hoegh-Guldberg O, Fine M, Skirving W, Johnstone R, Dove S, Strong A (2005) Coral bleaching following wintry weather. Limnol Oceanogr 50:265–271. https://doi.org/10.4319/lo.2005.50.1.0265

Hoegh-Guldberg O, Jacob D, Taylor M, Guillén Bolaños T, Bindi M, Brown S, Camilloni IA, Diedhiou A, Djalante R, Ebi K, Engelbrecht F, Guiot J, Hijioka Y, Mehrotra S, Hope CW, Payne AJ, Pörtner HO, Seneviratne SI, Thomas A, Warren R, Zhou G (2019) The human imperative of stabilizing global climate change at 1.5°C. Science (80-) 365:. https://doi.org/10.1126/science.aaw6974

Hoegh-Guldberg O, Kennedy E V., Beyer HL, McClennen C, Possingham HP (2018) Securing a Long-term Future for Coral Reefs. Trends Ecol Evol 33:936–944. https://doi.org/10.1016/j.tree.2018.09.006

Holbrook NJ, Scannell HA, Sen Gupta A, Benthuysen JA, Feng M, Oliver ECJ, Alexander L V., Burrows MT, Donat MG, Hobday AJ, Moore PJ, Perkins-Kirkpatrick SE, Smale DA, Straub SC, Wernberg T (2019) A global assessment of marine heatwaves and their drivers. Nat Commun 10:1–13. https://doi.org/10.1038/s41467-019-10206-z

Howells EJ, Abrego D, Meyer E, Kirk NL, Burt JA (2016) Host adaptation and unexpected symbiont partners enable reef-building corals to tolerate extreme temperatures. Glob Chang Biol 22:2702– 2714. https://doi.org/10.1111/gcb.13250

Hughes TP, Anderson KD, Connolly SR, Heron SF, Kerry JT, Lough JM, Baird AH, Baum JK, Berumen ML, Bridge TC, Claar DC, Eakin CM, Gilmour JP, Graham NAJ, Harrison H, Hobbs JPA, Hoey AS, Hoogenboom M, Lowe RJ, McCulloch MT, Pandolfi JM, Pratchett M, Schoepf V, Torda G, Wilson SK (2018) Spatial and temporal patterns of mass bleaching of corals in the Anthropocene. Science (80-) 359:80–83. https://doi.org/10.1126/science.aan8048

Hughes TP, Kerry JT, Álvarez-Noriega M, Álvarez-Romero JG, Anderson KD, Baird AH, Babcock RC, Beger M, Bellwood DR, Berkelmans R, Bridge TC, Butler IR, Byrne M, Cantin NE, Comeau S, Connolly SR, Cumming GS, Dalton SJ, Diaz-Pulido G, Eakin CM, Figueira WF, Gilmour JP, Harrison HB, Heron SF, Hoey AS, Hobbs JPA, Hoogenboom MO, Kennedy E V., Kuo CY, Lough JM, Lowe RJ, Liu G, McCulloch MT, Malcolm HA, McWilliam MJ, Pandolfi JM, Pears RJ, Pratchett MS, Schoepf V, Simpson T, Skirving WJ, Sommer B, Torda G, Wachenfeld DR, Willis BL, Wilson SK (2017) Global warming and recurrent mass bleaching of corals. Nature 543:373–377. https://doi.org/10.1038/nature21707

Hulbert SH (1984) Pseudoreplication and the Design of Ecological Field Experiments. Ecol Monogr 54:187–211

Iskin U, Filiz N, Cao Y, Neif ÉM, Öğlü B, Lauridsen TL, Davidson TA, Søndergaard M, Tavsanoğlu ÜN, Beklioğlu M, Jeppesen E (2020) Impact of nutrients, temperatures, and a heat wave on zooplankton community structure: An experimental approach. Water (Switzerland) 12:1–19. https://doi.org/10.3390/w12123416

Jiménez-Quiroz M del C, Cervantes-Duarte R, Funes-Rodríguez R, Barón-Campis SA, García-Romero F de J, Hernández-Trujillo S, Hernández-Becerril DU, González-Armas R, Martell-Dubois R, Cerdeira-Estrada S, Fernández-Méndez JI, González-Ania L V., Vásquez-Ortiz M, Barrón-Barraza FJ (2019) Impact of “The Blob” and “El Niño” in the SW Baja California Peninsula: Plankton and environmental variability of Bahia Magdalena. Front Mar Sci 6:1–23. https://doi.org/10.3389/fmars.2019.00025

Jokiel PL, Coles SL (1990) Response of Hawaiian and other Indo-Pacific reef corals to elevated temperature. Coral Reefs 8:155–162. https://doi.org/10.1007/BF00265006

Jones T, Parrish JK, Peterson WT, Bjorkstedt EP, Bond NA, Ballance LT, Bowes V, Hipfner JM, Burgess HK, Dolliver JE, Lindquist K, Lindsey J, Nevins HM, Robertson RR, Roletto J, Wilson L, Joyce T, Harvey J (2018) Massive Mortality of a Planktivorous Seabird in Response to a Marine Heatwave. Geophys Res Lett 45:3193–3202. https://doi.org/10.1002/2017GL076164

Kim SW, Sampayo EM, Sommer B, Sims CA, Gómez-Cabrera M del C, Dalton SJ, Beger M, Malcolm HA, Ferrari R, Fraser N, Figueira WF, Smith SDA, Heron SF, Baird AH, Byrne M, Eakin CM, Edgar R, Hughes TP, Kyriacou N, Liu G, Matis PA, Skirving WJ, Pandolfi JM (2019) Refugia under threat: mass bleaching of coral assemblages in high-latitude eastern Australia. Glob Chang Biol 00:1–14. https://doi.org/10.1111/gcb.14772

Kirk NL, Howells EJ, Abrego D, Burt JA, Meyer E (2018) Genomic and transcriptomic signals of thermal tolerance in heat-tolerant corals (Platygyra daedalea) of the Arabian/Persian Gulf. Mol Ecol 27:5180–5194. https://doi.org/10.1111/mec.14934

LaJeunesse TC, Parkinson JE, Gabrielson PW, Jeong HJ, Reimer JD, Voolstra CR, Santos SR (2018) Systematic Revision of Symbiodiniaceae Highlights the Antiquity and Diversity of Coral Endosymbionts. Curr Biol 28:2570–2580.e6. https://doi.org/10.1016/j.cub.2018.07.008

Liew YJ, Howells EJ, Wang X, Michell CT, Burt JA, Idaghdour Y, Aranda M (2020) Intergenerational epigenetic inheritance in reef-building corals. Nat Clim Chang 10:254–259. https://doi.org/10.1038/s41558-019-0687-2

Little RJ, Rubin DB (2002) Statistical analysis with missing data. John Wiley & Sons, Ltd

Liu G, Heron SF, Mark Eakin C, Muller-Karger FE, Vega-Rodriguez M, Guild LS, de la Cour JL, Geiger EF, Skirving WJ, Burgess TFR, Strong AE, Harris A, Maturi E, Ignatov A, Sapper J, Li J, Lynds S (2014) Reef-scale thermal stress monitoring of coral ecosystems: New 5-km global products from NOAA coral reef watch. Remote Sens 6:11579–11606. https://doi.org/10.3390/rs61111579

Liu G, Rauenzahn J, Heron S, Eakin C, Skirving W, Christensen T, Strong A, Li J (2013) NOAA Coral Reef Watch 50 km Satellite Sea Surface Temperature-Based Decision Support System for Coral Bleaching Management

Liu G, Strong AE, Skirving W (2003) Remote sensing of sea surface temperatures during 2002 barrier reef coral bleaching. Eos (Washington DC) 84:2002–2004. https://doi.org/10.1029/2003EO150001

Marshall PA, Baird AH (2000) Bleaching of corals on the Great Barrier Reef: Differential susceptibilities among taxa. Coral Reefs 19:155–163. https://doi.org/10.1007/s003380000086

Matz M V., Treml EA, Aglyamova G V., Bay LK (2018) Potential and limits for rapid genetic adaptation to warming in a Great Barrier Reef coral. PLoS Genet 14:1–19. https://doi.org/10.1371/journal.pgen.1007220

McClanahan TR, Darling ES, Maina JM, Muthiga NA, D’Agata S, Leblond J, Arthur R, Jupiter SD, Wilson SK, Mangubhai S, Ussi AM, Guillaume MMM, Humphries AT, Patankar V, Shedrawi G, Pagu J, Grimsditch G (2020) Highly variable taxa-specific coral bleaching responses to thermal stresses. Mar Ecol Prog Ser 648:135–151. https://doi.org/10.3354/meps13402

Mumby PJ, Chisholm JRM, Edwards AJ, Andrefouet S, Jaubert J (2001) Cloudy weather may have saved Society Island reef corals during the 1998 ENSO event. Mar Ecol Prog Ser 222:209–216. https://doi.org/10.3354/meps222209

Nakagawa S, Freckleton RP (2008) Missing inaction: the dangers of ignoring missing data. Trends Ecol Evol 23:592–596. https://doi.org/10.1016/j.tree.2008.06.014

Oliver ECJ, Donat MG, Burrows MT, Moore PJ, Smale DA, Alexander L V., Benthuysen JA, Feng M, Sen Gupta A, Hobday AJ, Holbrook NJ, Perkins-Kirkpatrick SE, Scannell HA, Straub SC, Wernberg T (2018) Longer and more frequent marine heatwaves over the past century. Nat Commun 9:1–12. https://doi.org/10.1038/s41467-018-03732-9

Piatt JF, Parrish JK, Renner HM, Schoen SK, Jones TT, Arimitsu ML, Kuletz KJ, Bodenstein B, García-Reyes M, Duerr RS, Corcoran RM, Kaler RSA, McChesney GJ, Golightly RT, Coletti HA, Suryan RM, Burgess HK, Lindsey J, Lindquist K, Warzybok PM, Jahncke J, Roletto J, Sydeman WJ (2020) Extreme mortality and reproductive failure of common murres resulting from the northeast Pacific marine heatwave of 2014-2016

Robin X, Turck N, Hainard A, Tiberti N, Lisacek F, Sanchez J-C, Müller M (2011) pROC: an open-source package for R and S+ to analyze and compare ROC curves. BMC Bioinformatics 12:. https://doi.org/10.1186/147121051277

Romero-Torres M, Acosta A, Palacio-Castro AM, Treml EA, Zapata FA, Paz-García DA, Porter JW (2020) Coral reef resilience to thermal stress in the Eastern Tropical Pacific. Glob Chang Biol 26:3880–3890. https://doi.org/10.1111/gcb.15126

Rue H, Martino S, Chopin N (2009) Approximate Bayesian inference for latent Gaussian models by using integrated nested Laplace approximations. J R Stat Soc Ser B Stat Methodol 71:319–392. https://doi.org/10.1111/j.1467-9868.2008.00700.x

Safaie A, Silbiger NJ, McClanahan TR, Pawlak G, Barshis DJ, Hench JL, Rogers JS, Williams GJ, Davis KA (2018) High frequency temperature variability reduces the risk of coral bleaching. Nat Commun 9:1–12. https://doi.org/10.1038/s41467-018-04074-2

Skirving W, Enríquez S, Hedley JD, Dove S, Eakin CM, Mason RAB, Cour JLD La, Liu G, Hoegh-Guldberg O, Strong AE, Mumby PJ, Iglesias-Prieto R (2018) Remote sensing of coral bleaching using temperature and light: Progress towards an operational algorithm. Remote Sens 10:p. https://doi.org/10.3390/rs10010018

Skirving W, Marsh B, De La Cour J, Liu G, Harris A, Maturi E, Geiger E, Mark Eakin C (2020) Coraltemp and the coral reef watch coral bleaching heat stress product suite version 3.1. Remote Sens 12:1–10. https://doi.org/10.3390/rs12233856

Skirving WJ, Heron SF, Marsh BL, Liu G, De La Cour JL, Geiger EF, Eakin CM (2019) The relentless march of mass coral bleaching: a global perspective of changing heat stress. Coral Reefs 38:547–557. https://doi.org/10.1007/s00338-019-01799-4

Smale DA, Wernberg T, Oliver ECJ, Thomsen M, Harvey BP, Straub SC, Burrows MT, Alexander L V., Benthuysen JA, Donat MG, Feng M, Hobday AJ, Holbrook NJ, Perkins-Kirkpatrick SE, Scannell HA, Sen Gupta A, Payne BL, Moore PJ (2019) Marine heatwaves threaten global biodiversity and the provision of ecosystem services. Nat Clim Chang 9:306–312. https://doi.org/10.1038/s41558-019-0412-1

Spady BL, Devotta DA, De La Cour J, Gomez AM, Morgan JA, Donner SD, Liu G, Skirving WJ, Vasile R, Geiger E, Marsh B, Eakin CM, Manzello DP (2021) Global Coral Bleaching Database (NCEI Accession 0228498). In: NOAA Natl. Centers Environ. Information. Unpubl. Dataset. https://www.ncei.noaa.gov/archive/accession/0228498. Accessed 1 Sep 2020

Sully S, Burkepile DE, Donovan MK, Hodgson G, van Woesik R (2019) A global analysis of coral bleaching over the past two decades. Nat Commun 10:1–5. https://doi.org/10.1038/s41467-019-09238-2

Ummenhofer CC, Meehl GA (2017) Extreme weather and climate events with ecological relevance: A review. Philos Trans R Soc B Biol Sci 372:. https://doi.org/10.1098/rstb.2016.0135

Veron JEN (1995) Corals in space and time: the biogeography and evolution of the Scleractinia. Cornell University Press

von Biela VR, Arimitsu ML, Piatt JF, Heflin BM, Schoen S (2019) Extreme reduction in condition of a key forage fish during the Pacific marine heatwave of 2014–2016. Mar Ecol Prog Ser 613:171–182

Weeks SJ, Anthony KRN, Bakun A, Feldman GC, Hoegh-Guldberg O (2008) Improved predictions of coral bleaching using seasonal baselines and higher spatial resolution. Limnol Oceanogr 53:1369–1375. https://doi.org/10.4319/lo.2008.53.4.1369

Wellington GM, Glynn PW, Strong AE, Navarrete SA, Wieters E, Hubbard D (2001) Crisis on coral reefs linked to climate change. Eos (Washington DC) 82:1–5. https://doi.org/10.1029/01EO00001

Wiedenmann J, D’Angelo C, Smith EG, Hunt AN, Legiret FE, Postle AD, Achterberg EP (2013) Nutrient enrichment can increase the susceptibility of reef corals to bleaching. Nat Clim Chang 3:160–164. https://doi.org/10.1038/nclimate1661

Wyatt ASJ, Leichter JJ, Toth LT, Miyajima T, Aronson RB, Nagata T (2019) Heat accumulation on coral reefs mitigated by internal waves. Nat Geosci. https://doi.org/10.1038/s41561-019-0486-4

Wyatt ASJ, Leichter JJ, Toth LT, Miyajima T, Aronson RB, Nagata T (2020) Heat accumulation on coral reefs mitigated by internal waves. Nat Geosci 13:28–34. https://doi.org/10.1038/s41561-019-0486-4

Zuur AF, Ieno EN (2017) Beginner’s Guide to Spatial, Temporal and Spatial-Temporal Ecological Data Analysis with R-INLA. Volume II GAM and Zero-Inflated Models. Highland Statistics Ltd, 1

