## Supplementary Materials for "Fine-tuning heat stress algorithms to optimise global predictions of mass coral bleaching"

1. Coral Bleaching Data Filtration
2. Model Validation
3. Patchiness Simulation Test
4. Best-performing Spatiotemporal GLM example - Estimated Posterior Distributions
5. Best-performing Spatiotemporal GLM example - Spatiotemporal Correlation
6. References

1. Coral Bleaching Data Filtration

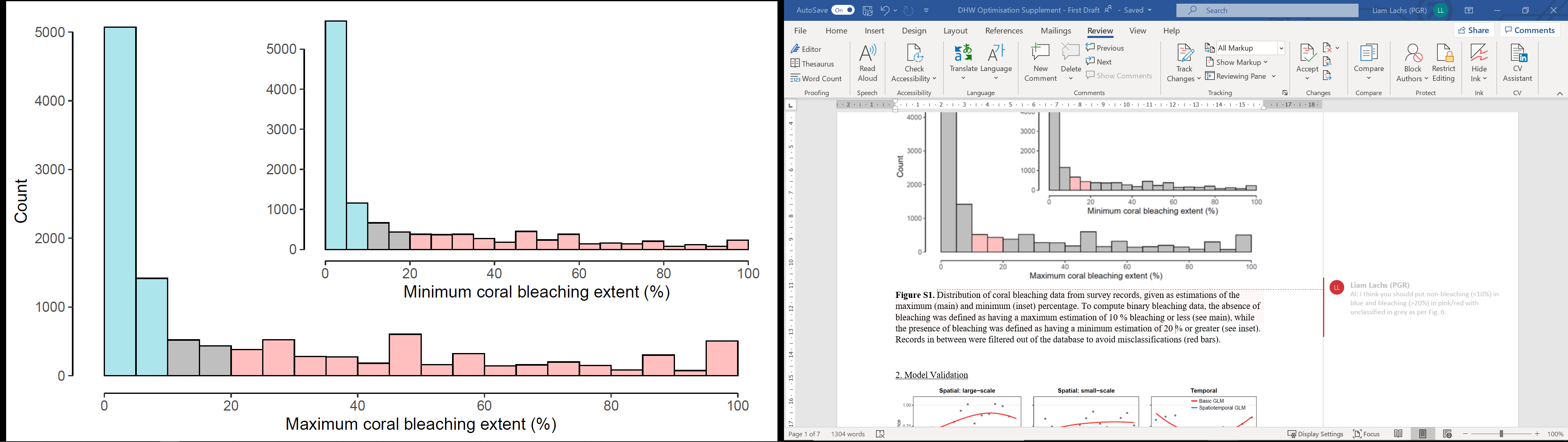

**Figure S1.** Distribution of coral bleaching data from survey records, given as estimations of the maximum (main) and minimum (inset) percentage. To compute binary bleaching data, the absence of bleaching was defined as having a maximum estimation of 10 % bleaching or less (see main, blue bars), while the presence of bleaching was defined as having a minimum estimation of 20 % or greater (see inset, red bars). Records in between were filtered out of the database to avoid misclassifications (grey bars).

2. Model Validation

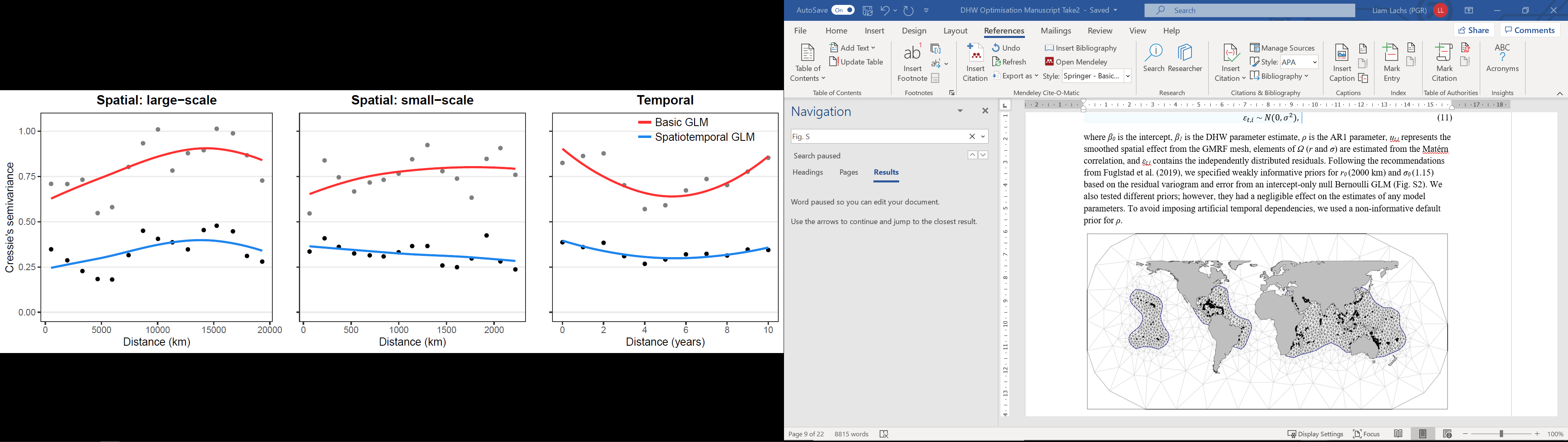

**Figure S2.** Comparison of residual correlation (large-scale spatial, small-scale spatial, and temporal) for a basic Generalised Linear Model (GLM) (red) and a spatiotemporal GLM (blue). Both GLMs were fit in the form: Bleaching ~ DHW_op_. Note lower spatial and temporal autocorrelation in model residuals for the spatiotemporal GLM.

3. Patchiness Simulation Test

*Aims*

It is well-known that patchy data can cause biased estimates of model parameters, for example, the slope, intercept and standard error of a linear regression (Bihrmann and Ersbøll 2015). These biases can also extend to model comparison criteria such as Akaike Information Criterion (AIC) or Deviance Information Criterion (DIC) used in this study (Nakagawa and Freckleton 2008). Although we employed the most extensive global coral bleaching dataset available, observations were not distributed consistently through space and time. Observations were limited to the location of coral reefs, with more observations in recent years and on the more accessible reefs, and no observations for some geographical regions in certain years (mostly due to an absence of bleaching). Therefore, to validate this study of optimising heat stress metrics using Bayesian Generalised Linear Models (GLMs), we also needed to address model biases relating to patchy data, or as we refer to, patchiness.

*Methods*

To address potential bias due to patchiness in the coral bleaching dataset, we have performed a simulation test across four patchiness scenarios: a regular grid, a spatially patchy grid, a spatiotemporally patchy grid, and the true level of patchiness (i.e. actual bleaching survey sites). The overall aim was to assess whether patchiness affected the outcome of the Bayesian GLMs presented in this study. This simulation test was performed for each patchiness scenario independently, with the same methodology. For clarity, we describe this step-by-step workflow for the regular-grid scenario only, aided by a conceptual diagram (Fig. S1):

1. An INLA GLM (the “true model”) was fitted to bleaching observations based on one DHW_test_ metric (threshold = MMM + 1 ⁰C, window: 12-weeks) and the spatiotemporal random effect. The same model settings were applied here, as in the main manuscript models, except a coarser mesh was defined to improve runtime (maximum triangle edge length = 4000km, convex hull = -0.06). This translates to 193 nodes per timestep, which exceeds the recommended minimum value of 100 nodes per timestep (Bakka et al. 2018).
2. To facilitate the simulations, 1000 sets of parameters (*β_0_, β_1_, Ω, ρ*) were randomly sampled from their posterior distributions in the “true model”.
3. Aside to this, a regular grid of points bounding all observations in the study area was defined (250 × 250, N = 62,500). DHW values from observed coordinates in each year were interpolated to the regular-gridded coordinates in each corresponding year using nearest neighbour interpolation in R (gstat library), where a maximum of 1000 neighbours are used for interpolations at any given point.
4. Using each of the 1000 sets of parameters from Step 2, 1000 sets of bleaching data were simulated from the regular DHW grid. This was achieved by feeding all relevant parameters into equation 8 in the original manuscript to get values of *π_t,i_*. Then following equation 5 in the original manuscript (rbinom function in R) values of *π_t,i_* are then converted to Bernoulli draws (1 or 0, i.e. the simulated bleaching dataset). Given that each of the 1000 simulated bleaching dataset were based on the same interpolated DHW values, the only variations among them were due to the random sampling of model parameter values (*β_0_, β_1_, Ω, ρ*).
5. New INLA GLMs (or “test models”) were fitted to the 1000 simulated datasets independently, as in Step 1. This produced a set of new parameter estimates (*β_0_, β_1_, Ω, ρ*) for each “test model”.
6. The posterior means of model parameters were then compared between the “true model” (Fig. S1, blue diamond) and the “test models” (Fig. S1, histogram and 95% CI). If patchiness is not a computational issue for the simulated dataset then the “true model” parameter should lie within the 95% confidence intervals of the 1000 “test model” parameters. Since this study is focused on bleaching and heat stress, we evaluated each simulation using *β_1_*, the DHW parameter.

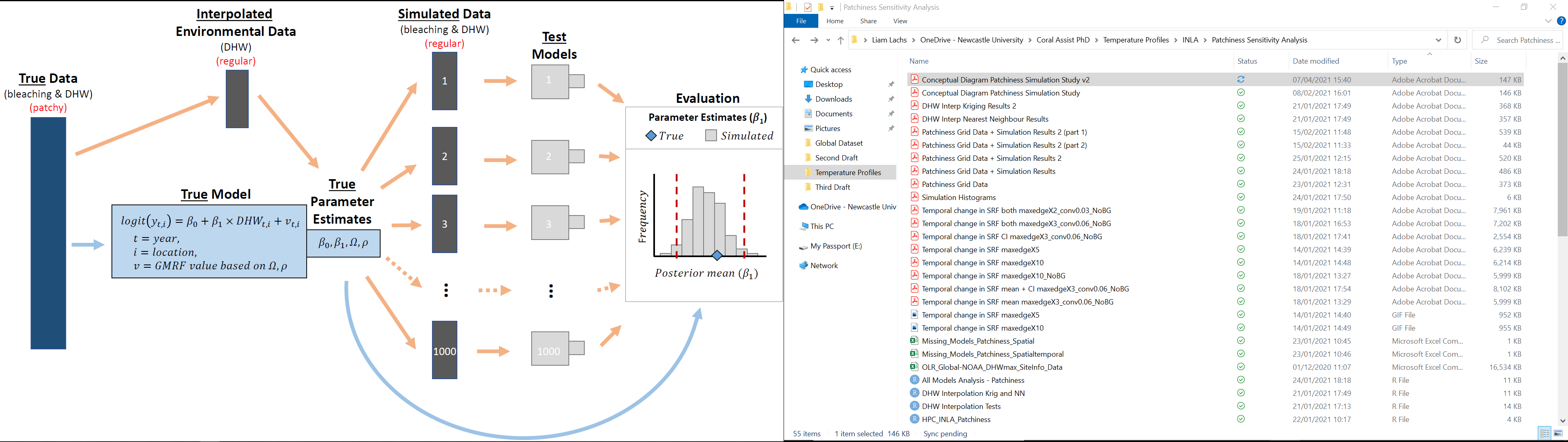
**Figure S3.** Conceptual workflow of the patchiness simulation test. A “true model” (spatiotemporal Generalised Linear Model in INLA) is built on observed bleaching and DHW data (blue colours), and is used to estimate posterior distributions for each model parameter (*β_0_, β_1_, Ω, ρ*). By combining 1000 sets of “true model” parameters with a regular grid of interpolated DHWs, corresponding bleaching values can be simulated for each interpolated DHW dataset (dark grey colour). Then “test models” can be fitted to each of these simulated datasets (light grey colour). The posterior means of estimated model parameters are then compared between the “true model” (blue diamond) and the “test models” (histogram and 95% confidence interval).

The other three patchiness scenarios (spatial, spatiotemporal, and true) were tested in the same way as described above with alterations to Step 3 (i.e., how the DHW grid was formed). The spatially patchy grid was derived from the regular grid, whereby all DHW datapoints were removed from areas that had no true observations (Fig. S2, West African coast, South America, and other points on land, etc.). The spatiotemporally patchy grid was derived from the spatially patchy grid, whereby DHW datapoints from each year were randomly removed following the temporal distribution of bleaching observations (Fig. S2). Finally, the true patchiness scenario utilised the coordinates of bleaching observations and true DHW values (Fig. S2).

*Results and Conclusion*

For all four scenarios of patchiness, the *β_1_* parameter estimate from the “true model” (c.f. Step 1) is within the 95% confidence intervals of *β_1_* parameter estimates from “test models” (Fig. S3). This shows that patchiness did not have an impact on R-INLA’s computational ability to estimate model parameters. Based on this fact, we conclude that the level of patchiness seen in this bleaching dataset is within the levels accommodated by R-INLA. This validates the assumption that there are no model biases based on the levels of patchiness (data gaps) seen in this dataset.

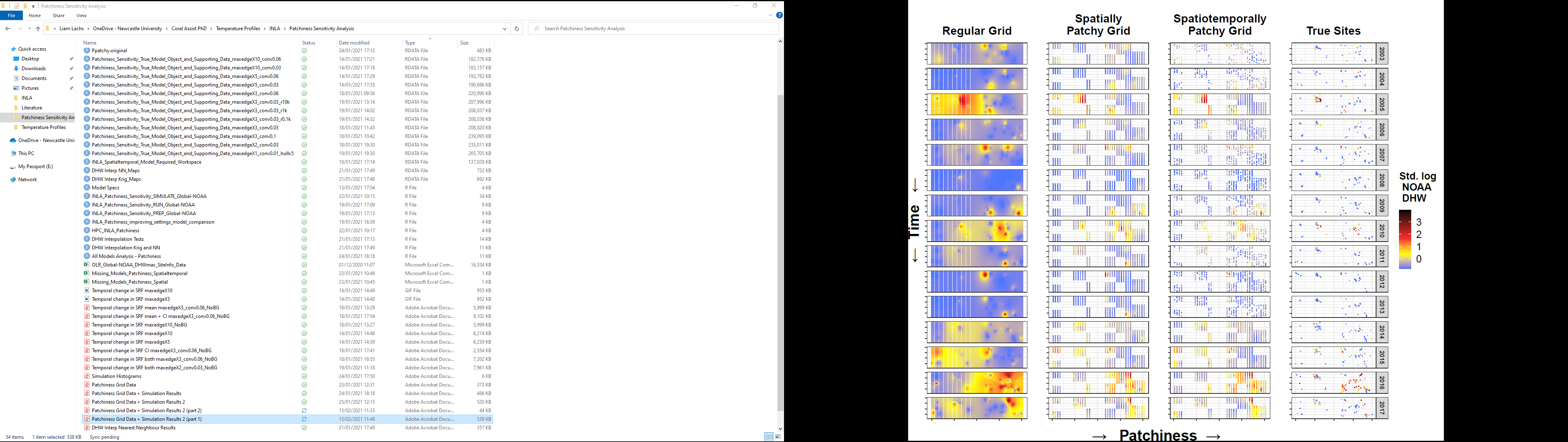

**Figure S4.** Dataset structure across a gradient of patchiness with four distinct scenarios: a regular grid of interpolated DHW values (left), a spatially patchy grid (middle left), a spatiotemporally patchy grid (middle right), and the true patchiness scenario with true DHW values (right). Each facet represents the same bounding box on a world map that surrounds all bleaching observations. Data is shown for each year from 2003 to 2017.

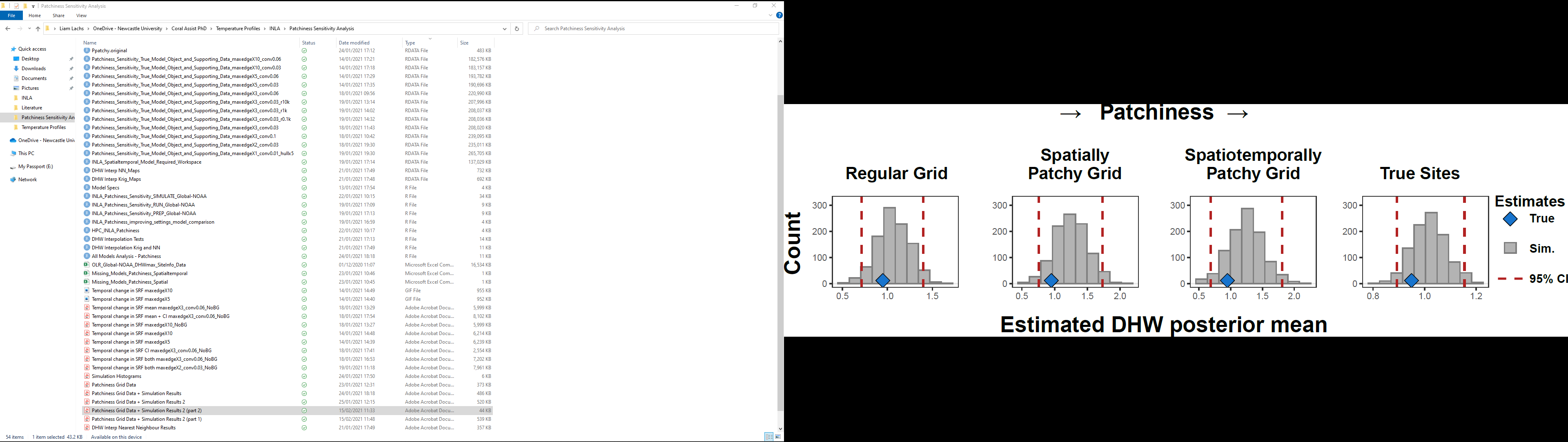

**Figure S5.** The results of the patchiness simulation test are shown for each patchiness scenario (as in Fig. S4), based on the mean of the *β_1_* parameter posterior distribution (i.e., DHW parameter). In each case, the “true” *β_1_* parameter estimate (blue diamonds) is within the 95% confidence intervals of the “test” *β_1_* parameter estimates (histograms).

4. Best-performing Spatiotemporal GLM example - Estimated Posterior Distributions

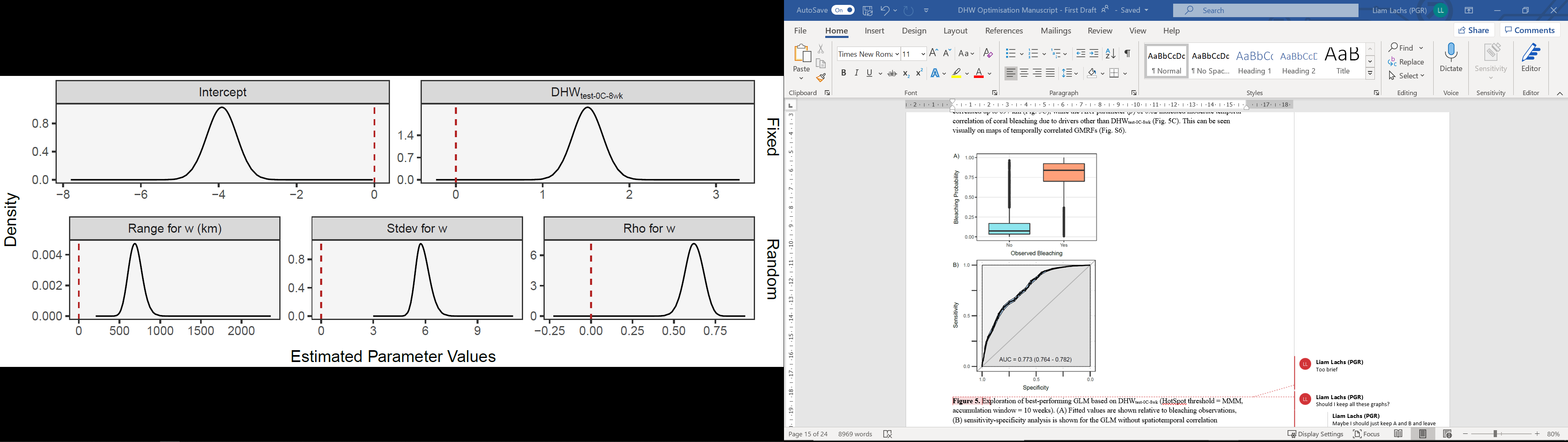

**Figure S6.** Posterior distributions of model parameters for one of the best performing Generalised Linear Models (Bleaching ~ DHW_test-0C-8wk_ using HotSpot threshold of MMM + 0 ⁰C and accumulation window of 8 week). Fixed effects (upper) and random effects (lower – spatiotemporal correlation) are shown. There is a clear positive association between heat stress (DHW_test-0C-8wk_) and coral bleaching. The range up to which spatial correlated random effect are correlated is approximately 600 km (i.e. drivers other than heat stress). Moderate positive temporal correlation is apparent from the rho value (AR1 parameter) of 0.62, meaning model error in one year is likely to be similar to the previous year.

5. Best-performing Spatiotemporal GLM example - Spatiotemporal Correlation

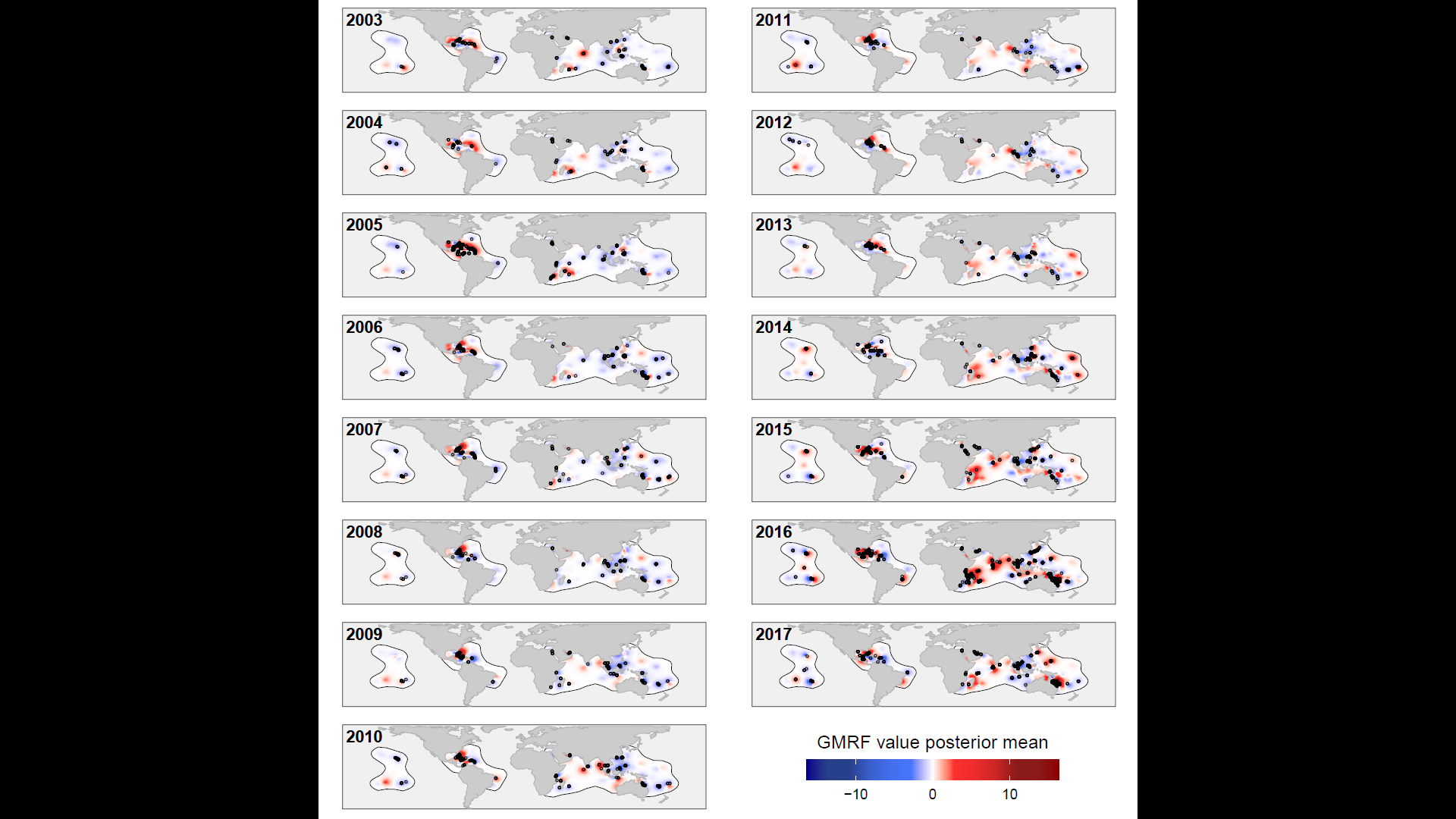

**Figure S7.** Gaussian Markov random fields through time for one of the best performing Generalised Linear Models (Bleaching ~ DHW_test-0C-8wk_ using HotSpot threshold of MMM + 0 ⁰C and accumulation window of 8 week). GMRF values (*v*_t,i_) show the strength of the spatiotemporally correlated random effect (i.e. drivers of bleaching other than DHW) at location *i* in year *t*.

6. References

Bakka H, Rue H, Fuglstad GA, Riebler A, Bolin D, Illian J, Krainski E, Simpson D, Lindgren F (2018) Spatial modeling with R-INLA: A review. Wiley Interdiscip Rev Comput Stat 10:1–24 . https://doi.org/10.1002/wics.1443

Bihrmann K, Ersbøll AK (2015) Estimating range of influence in case of missing spatial data: A simulation study on binary data. Int J Health Geogr 14: . https://doi.org/10.1186/1476-072X-14-1

Nakagawa S, Freckleton RP (2008) Missing inaction: the dangers of ignoring missing data. Trends Ecol Evol 23:592–596 . https://doi.org/10.1016/j.tree.2008.06.014
